# Top-down, knowledge-based genetic reduction of yeast central carbon metabolism

**DOI:** 10.1101/2021.08.24.457526

**Authors:** Eline D. Postma, Lucas G.F. Couwenberg, Roderick N. van Roosmalen, Jordi Geelhoed, Philip A. de Groot, Pascale Daran-Lapujade

## Abstract

*Saccharomyces cerevisiae*, whose evolutionary past includes a whole-genome duplication event, is characterised by a mosaic genome configuration with substantial apparent genetic redundancy. This apparent redundancy raises questions about the evolutionary driving force for genomic fixation of ‘minor’ paralogs and complicates modular and combinatorial metabolic engineering strategies. While isoenzymes might be important in specific environments, they could be dispensable in controlled laboratory or industrial contexts. The present study explores the extent to which the genetic complexity of the central carbon metabolism (CCM) in *S. cerevisiae*, here defined as the combination of glycolysis, pentose phosphate pathway, tricarboxylic acid cycle and a limited number of related pathways and reactions, can be reduced by elimination of (iso)enzymes without major negative impacts on strain physiology. Cas9-mediated, groupwise deletion of 35 from the 111 genes yielded a ‘minimal CCM’ strain, which despite the elimination of 32 % of CCM-related proteins, showed only a minimal change in phenotype on glucose-containing synthetic medium in controlled bioreactor cultures relative to a congenic reference strain. Analysis under a wide range of other growth and stress conditions revealed remarkably few phenotypic changes of the reduction of genetic complexity. Still, a well-documented context-dependent role of *GPD1* in osmotolerance was confirmed. The minimal CCM strain provides a model system for further research into genetic redundancy of yeast genes and a platform for strategies aimed at large-scale, combinatorial remodelling of yeast CCM.

## Introduction

The fundamental challenge of defining the minimum complement of genes required for life has been addressed by theoretical as well as experimental approaches. Bottom-up and top-down strategies mainly focused on bacteria with small genomes (Hashimoto *et al*. 2005; Hutchison *et al*. 2016; Komatsu *et al*. 2010; Lara and Gosset 2019; Lieder *et al*. 2015; Morimoto *et al*. 2008; Mutschler *et al*. 2019; Suzuki *et al*. 2005; Zhu *et al*. 2017). Their larger genome sizes might appear to make eukaryotic microorganisms less relevant for this type of research. However, they do offer attractive models to explore the biological significance of (apparent) genetic redundancy. Different evolutionary advantages have been proposed for the fixation of duplicated genes in genomes, including provision of a molecular landscape for functional (minor or major) innovation (e.g., neo- and subfunctionalization), a functional backup, gene dosage effects or increased buffering to respond to environmental cues (Conant and Wolfe 2008; Escalera-Fanjul *et al*. 2019). Systematically identifying the physiological significance underlying gene fixation presents a daunting challenge.

With its relatively small genome (12 Mb), tractability and high genetic accessibility, the yeast *Saccharomyces cerevisiae* is a valuable model for fundamental research on minimal genetic requirements. *S. cerevisiae* underwent a whole genome duplication (WGD) approximately 100 million years ago as well as Smaller-Scale Duplication (SSD) events. While 90 % of the WGD genes were lost during evolution, some duplicates remain (Escalera-Fanjul *et al*. 2019). As in humans, a substantial fraction of the total gene duplicates in *S. cerevisiae* originates from the WGD (approximately 63 % of duplicates in *S. cerevisiae* genome and 62 % in the human genome), while a smaller fraction originates from SSD (approximately 37 % of duplicates in *S. cerevisiae* genome and 38 % in the human genome) (Acharya and Ghosh 2016; Fares *et al*. 2013). While systematic, large scale studies like the construction of the yeast deletion collections (Costanzo *et al*. 2016; Giaever *et al*. 2002; Kuzmin *et al*. 2018; Winzeler *et al*. 1999), the synthetic genetic array projects (Costanzo *et al*. 2010; Tong *et al*. 2001; Tong *et al*. 2004) or the recent SCRaMbLE-based genome compaction (Luo *et al*. 2021), have provided valuable information on the dispensability of (a combination of) genes, the physiological role of many of these paralogous genes remains poorly defined.

In addition to the fundamental scientific questions raised by genetic redundancy, it also complicates genome engineering of *S. cerevisiae*. The conversion of substrate into product via native or engineered pathways, relies on the microbial host core pathways for the supply of metabolic precursors, energy-rich molecules and redox equivalents. These biochemical reactions are catalysed by sets of ‘metabolic’ genes that are characterized by a high genetic redundancy in eukaryotes (Escalera-Fanjul *et al*. 2019). Not only has the physiological role of many paralogous genes not been fully elucidated, but the manipulation of specific biochemical reactions is hindered by the presence of multiple paralogous genes with poorly known functions that are scattered over the 12 Mb, mosaic yeast genome and its 16 chromosomes. Additionally, expression of these redundant genes dissipates cellular resources (e.g., carbon, energy) that might be better invested in industrially-relevant properties such as high product yield or cellular robustness to the stressful environment of large-scale fermentation.

To tackle these fundamental and applied challenges, taking glycolysis and ethanolic fermentation as starting point, Solis-Escalante and colleagues pioneered the genetic reduction of central carbon metabolism in *S. cerevisiae*. The set of 26 genes encoding the (iso)enzymes catalysing 12 reactions was reduced to 13 genes (Solis-Escalante *et al*. 2015). Remarkably, this 50 % genetic reduction did not result in any visible phenotypic effect, although a wide range of growth conditions was tested. These observations argued against gene dosage being a strong driving force in the evolution of Crabtree positive yeasts (Escalera-Fanjul *et al*. 2019; Solis-Escalante *et al*. 2015) and raised questions on the mechanisms involved the fixation of these gene duplicates in the *S. cerevisiae* genome. A recent study suggests that the role of the redundant paralogs might be highly context-dependent and that some relevant conditions were not tested by Solis-Escalante *et al*. (2015) (e.g. the role of Pyruvate kinase 2 in the utilization of three-carbon substrates such as dihydroxyacetone (Bradley *et al*. 2019)). The surprising lack of phenotype of the “Minimal Glycolysis” yeast strain (called MG) enabled the construction of a genetically simplified version of the glycolytic pathway, which was subsequently relocalized to a single chromosomal locus (Kuijpers *et al*. 2016). The resulting yeast strain with a single-locus glycolysis presents a powerful tool to remodel the glycolytic pathway in two single steps into any redesigned (heterologous) version.

Glycolysis is an important but small part of Central Carbon Metabolism (CCM), a set of reactions required for the conversion of carbon feedstocks into any industrially-relevant product (Fig. 1). For cells, CCM is primarily the set of reactions that convert carbon sources into the 12 building blocks required for the synthesis of cellular components (Nielsen and Keasling 2016). CCM encompasses ca. 111 genes, with 66 % of duplicates (Fig. 2). Reducing its genetic complexity would be the first step in an attempt to construct a modular, designer yeast genome, with a single-locus CCM, as previously achieved for glycolysis and fermentation. Modular, specialised synthetic chromosomes could be ideal platforms for the centralisation of the CCM genes (Postma *et al*. 2021).

**Figure 1.**
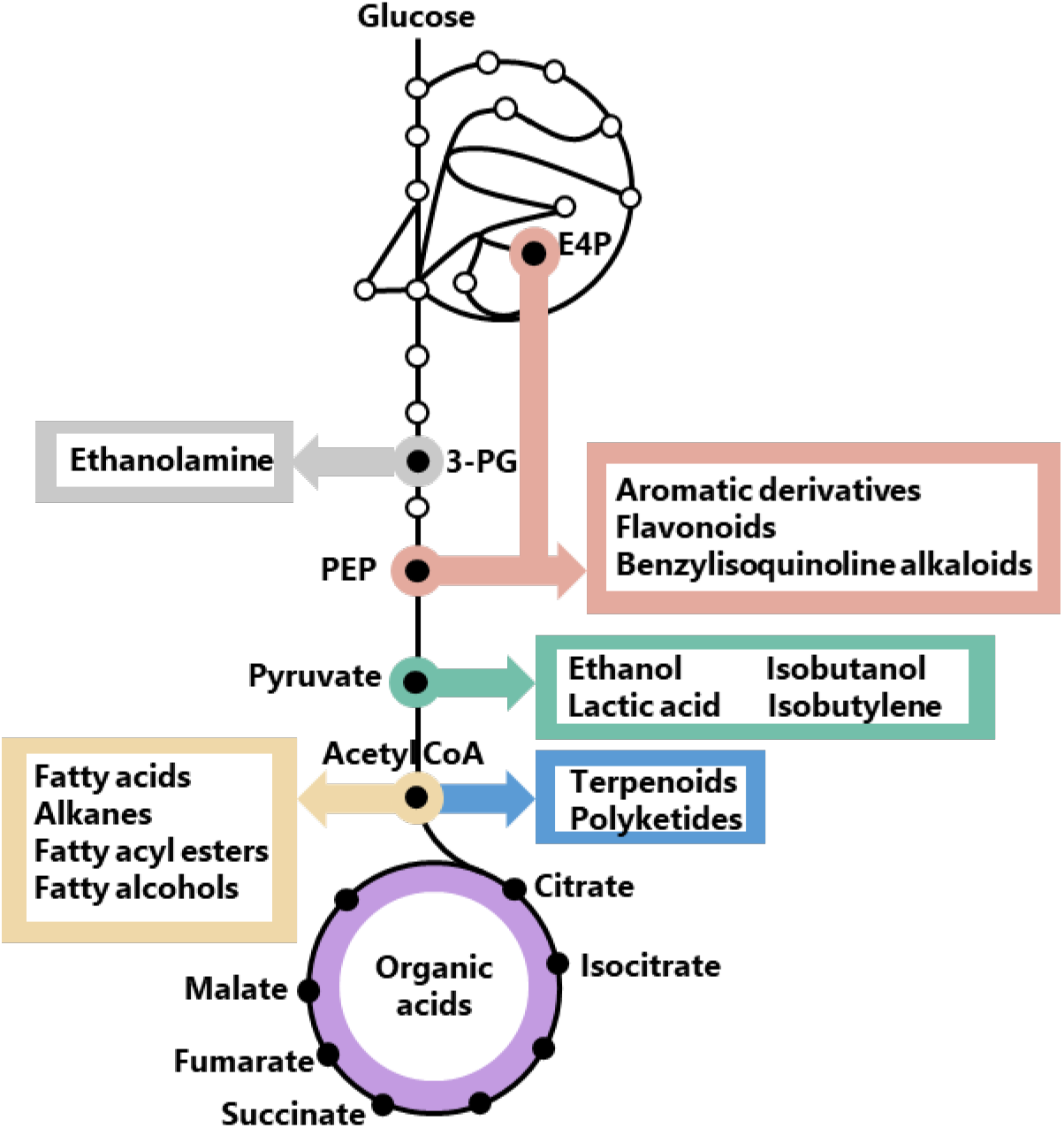
CCM precursors of industrially relevant chemicals. Abbreviations: 3-PG is 3-Phosphoglycerate and PEP is phosphoenolpyruvate.

**Figure 2.**
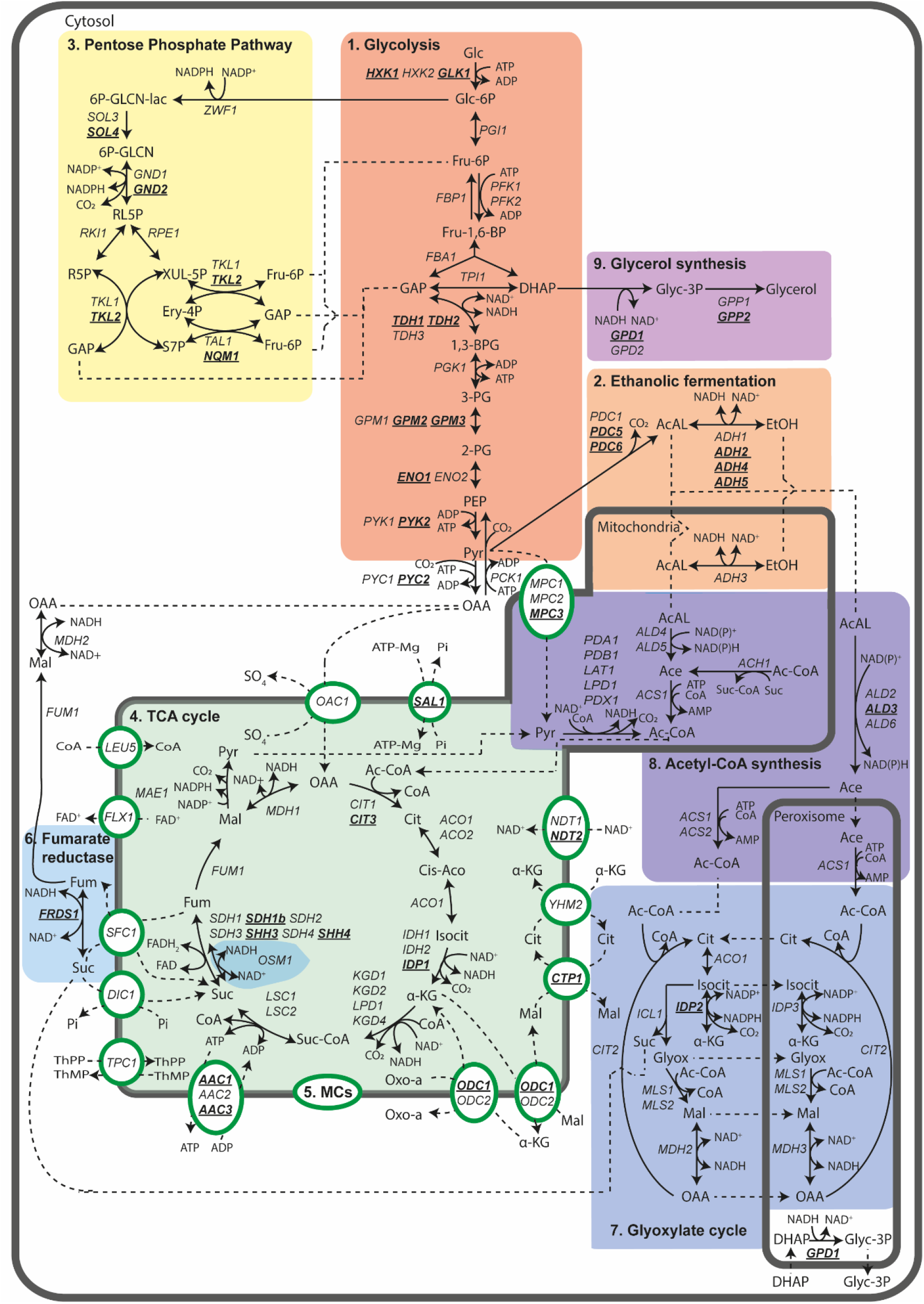
Reactions of the central carbon metabolism (CCM) of *S. cerevisiae* considered for genetic reduction. Schematic representation of sub-processes and pathways in CCM metabolic pathways considered for genetic reduction in this study. Enzyme-catalysed conversions (black lines) and transport processes (dotted lines) are shown between intermediates and through mitochondrial transporters (circles and ovals), respectively. Directionality and reversibility of reactions was based on https://pathway.yeastgenome.org/. Enzyme localization was based on literature information. Genes retained in the genetic reduction strategy are shown in black, genes selected for deletion in the minimal CCM strain are indicated in bold and underlined. Occurrence of pathways in different cellular compartments is shown by grey borders. Simplifications have been made for visualization reasons, for example H_2_O and inorganic phosphate are not shown. Abbreviations can be found in the abbreviation list.

The main goal of this study was to explore the extent to which the number of genes encoding CCM enzymes in *S. cerevisiae* can be reduced without substantially affecting fitness under a set of chosen growth conditions. To this end, redundancies were first predicted based on literature data on gene expression, enzyme activities and phenotype of (single) deletion mutants. Subsequently, phenotypes of mutants with mutations in sets of genes encoding CCM enzymes were tested under a wider range of growth conditions. In this first attempt of genetic reduction of yeast CCM at this scale, special attention is given to possible synergistic effects of mutations that were previously studied in separate strains.

## Results

### Genetic reduction strategy

In this study the CCM of *S. cerevisiae* was defined as the set of biochemical reactions encompassed by glycolysis, ethanolic fermentation, pentose-phosphate pathway, acetyl-CoA synthesis, tricarboxylic acid cycle, anaplerosis, gluconeogenesis, glyoxylate cycle and glycerol metabolism. As CCM reactions occur in multiple compartments, mitochondrial transporters were also considered (Fig. 2). Transport over the peroxisomal membrane was not considered as this phenomenon is poorly studied (DeLoache *et al*. 2016). For the construction of a minimal CCM stain, decisions to remove or retain genes were based on (i) transcript levels from an expression compendium encompassing 170 different cultivation conditions (Knijnenburg *et al*. 2009), (ii) enzyme activities in cell extracts of mutant strains when data were available, and (iii) on reported phenotypes of null mutants. Genes encoding proteins with reported secondary (‘moonlighting’) functions or proteins known to cause auxotrophy upon deletion were retained (Gancedo and Flores 2008).

Genes were classified as functionally redundant when at least 75 % of the specific growth rate of the congenic reference strains CEN.PK113-7D (Ura^+^) or IMX581 (Ura^-^) was retained during aerobic batch cultivation on synthetic medium supplied with either glucose or ethanol. Ethanol-grown cultures were included as, in contrast to glucose, ethanol can only be dissimilated by respiration and because its metabolism involves different sets of CCM enzymes and transporters. In addition, testing for growth on ethanol ensured that the intensive engineering undergone by the strains, including removal of several mitochondrial proteins, did not cause respiratory deficiency.

The previously constructed MG strain, in which 13 out of the 26 existing paralogs of genes encoding glycolytic enzymes and fermentation enzymes were deleted without the detection of major phenotypes, was used as starting point of the present CCM reduction endeavour. To identify any synergistic effects between the newly introduced deletions and the 13 deletions already present in the MG strain, a congenic naïve reference strain (IMX581) with a full complement of glycolytic and fermentation genes was also used in parallel to MG for serial deletions. To accelerate the deletion workflow, genes involved in individual pathways or processes were deleted in sets of two to four (Fig. 3). When substantial loss of fitness was observed, the contribution of individual deletions was dissected by constructing additional strains with various combinations and numbers of deletions.

**Figure 3.**
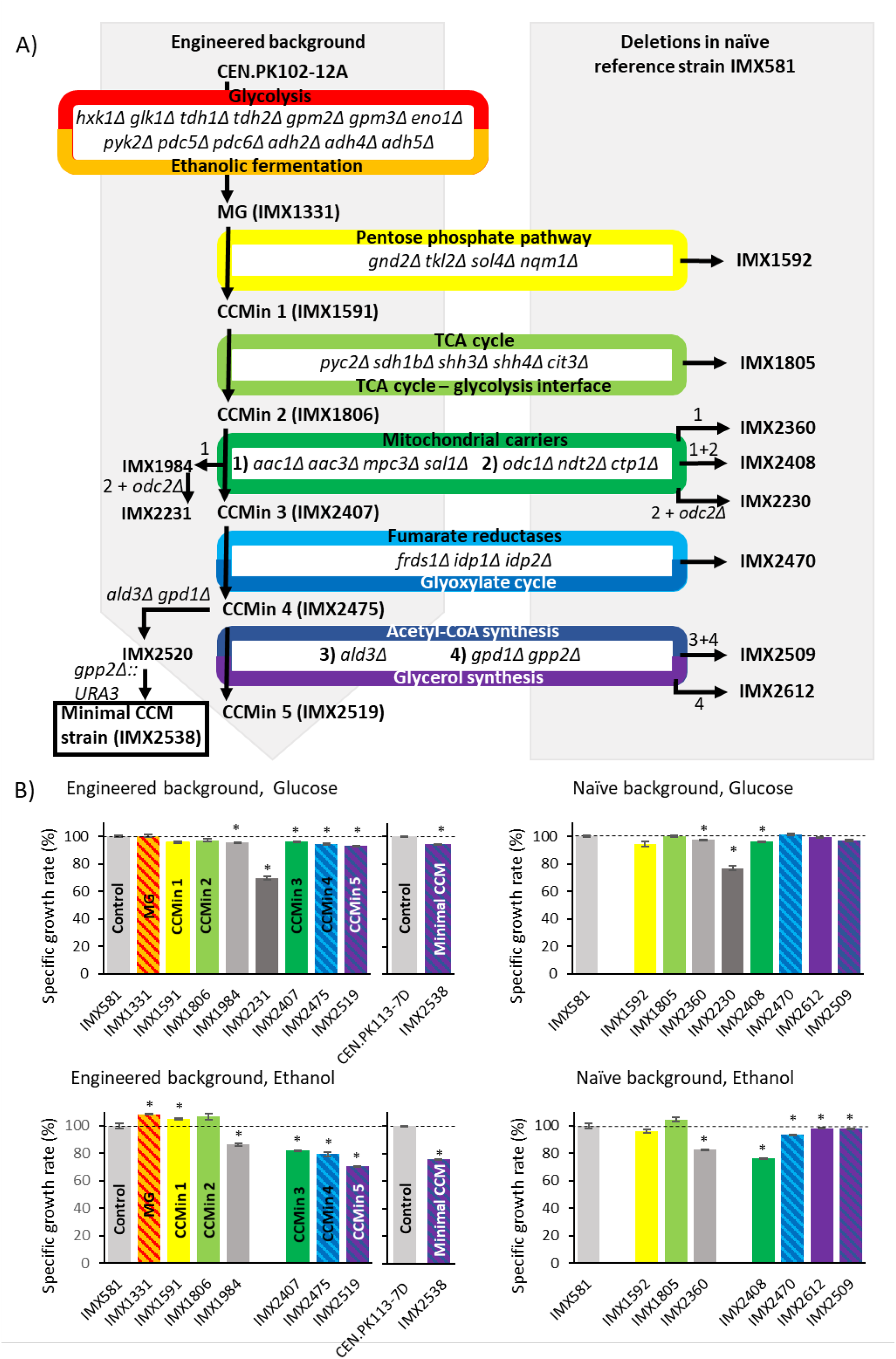
Deletion strategy and specific growth rates of resulting strains. Construction workflow for relevant *S. cerevisiae* strains (A) and their respective specific growth rates measured in shake flask rate on synthetic medium with glucose (SMD) or ethanol (SME) as carbon source, supplemented with uracil (B). Specific growth rates represent average and standard deviation of measurements on independent duplicate cultures for each strain and are expressed as a percentage of specific growth rate of the naïve uracil auxotrophic reference strain *S. cerevisiae* IMX581 or the naïve uracil prototrophic reference strain *S. cerevisiae* CEN.PK113-7D. Significant differences in specific growth rate relative to the control strain are indicated with a * (two-tailed paired homoscedastic t-test p<0.05).

### Deletion of 35 CCM genes has minimal impact on specific growth rate on chemically defined glucose medium

#### Pentose-phosphate pathway

The pentose-phosphate pathway (PPP) reduces cellular NADP^+^, generates ribose-5-phosphate and erythrose-4-phosphate for nucleic acid and amino-acid synthesis and, in strains engineered for pentose fermentation, acts as dissimilatory pathway (Jansen *et al*. 2017).

Four of the seven reactions in the PPP are catalyzed by pairs of isoenzymes encoded by WGD paralogs, for which sequence similarities range from 47 % (*SOL3* and *SOL4*) to 87 % (*GND1* and *GND2*) (Suppl. Table S1). Based on transcript levels across a wide ranges of cultivation conditions (Knijnenburg *et al*. 2009), *SOL4, GND2, TKL2* and *NQM1* were considered as minor paralogs. Moreover, deletion of *TKL2* and *NQM1* was previously reported not to affect growth on glucose synthetic medium (Lobo and Maitra 1982; Matsushika *et al*. 2012; Schaaff-Gerstenschläger *et al*. 1993). While similar *in vitro* enzyme activities were reported for Sol3 and Sol4 in cell extracts (Stanford *et al*. 2004), *SOL4* was deleted based on its consistently lower transcript level (Knijnenburg *et al*. 2009). Simultaneous deletion of *GND2, TKL2, SOL4* and *NQM1* in the naïve reference strain or in the MG strain, while retaining *SOL3*, *GND1*, *TKL1* and *TAL1*, did not significantly affect growth rate on either synthetic medium with 2 % (w/v) glucose (SMD) or 2 % (v/v) ethanol (SME) as carbon sources (strains IMX1592 (ppp^min^) and IMX1591 (called CCMin1: glyc^min^ fer^min^ ppp^min^), Fig. 3). Absence of a previously reported extended lag phase and slower growth on ethanol of *sol4* null mutants (Castelli *et al*. 2011) was not observed. This difference may be related to the use of different *S. cerevisiae* strain backgrounds.

#### Tricarboxylic acid cycle, anaplerotic reactions and gluconeogenesis

In addition to its dissimilatory role in oxidizing acetyl-CoA units to CO_2_, the tricarboxylic acid (TCA) cycle supplies precursors, NADH, FADH_2_ and ATP (Barnett 2003). During growth on fermentable sugars, the TCA cycle is a mitochondrial pathway, with acetyl-CoA resulting from oxidative decarboxylation of pyruvate by the pyruvate dehydrogenase (PDH) complex. To replenish use of TCA-cycle intermediates for biosynthesis, the cycle’s acceptor molecule oxaloacetate can be imported from the cytosol, where it is produced by carboxylation of pyruvate. The nine biochemical reactions of the TCA cycle involve 22 mitochondrial enzymes, which show little genetic redundancy. Two reactions are catalysed by single enzymes, Mdh1 and Fum1, while three steps are catalysed by complexes of two to five proteins. Deletion of genes encoding individual subunits of the α-ketoglutarate and succinyl-CoA synthetase complexes renders the complexes dysfunctional (Cupp and McAlister-Henn 1991; Cupp and McAlister-Henn 1992; Dickinson *et al*. 1986; Przybyla-Zawislak *et al*. 1998; Repetto and Tzagoloff 1989; Repetto and Tzagoloff 1990). In contrast, the succinate dehydrogenase complex, in which four functions are performed by seven proteins, does show some redundancy. *SDH1*, *SDH3* and *SDH4* have *SDH1b*, *SHH3*, and *SHH4*, respectively, as homologs originating from the WGD, while *SDH2* is a unique gene (Bullis and Lemire 1994; Chapman *et al*. 1992; Colby *et al*. 1998; Gebert *et al*. 2011; Lombardo *et al*. 1990; Szeto *et al*. 2012). Deletion of *SDH1b, SHH3* and *SHH4* has minor or no effect on complex integrity and yeast physiology and these genes are considered as functionally redundant (Chang *et al*. 2015; Colby *et al*. 1998; Szeto *et al*. 2012). Citrate synthase (Cit1 and Cit3) and isocitrate dehydrogenase (Idh1, Idh2 and Idp1) have functionally redundant mitochondrial enzymes. Based on expression data and lack of a phenotypic difference during fermentative and respiratory growth, Cit3 and the NADP^+^-dependent Idp1 were considered to be redundant (Haselbeck *et al*. 1992; Jia *et al*. 1997; Knijnenburg *et al*. 2009; Zhao and McAlister-Henn 1996). Single deletion of *ACO1* or *ACO2*, which encode aconitase isoenzymes, causes amino acid auxotrophies (Fazius *et al*. 2012; Gangloff *et al*. 1990). Idh1 and Idh2 are part of a complex and are both required for isocitrate dehydrogenase activity (Cupp and McAlister-Henn 1991; Cupp and McAlister-Henn 1992). Based on this information, only five of the 22 TCA cycle mitochondrial proteins were considered functionally redundant and, therefore, selected as candidates for elimination: Cit3, Idp1, Sdh1b, Shh3 and Shh4. *IDP1* was targeted in a later deletion round, along with extra-mitochondrial paralogs of TCA-cycle that are part of the glyoxylate cycle.

There are several enzymes that form an interface between the TCA cycle and glycolysis. The WGD paralog pair *PYC1* and *PYC2* encodes isoenzymes of the anaplerotic enzyme pyruvate carboxylase. Transcript levels of these two highly similar genes (92 %, Suppl. Table S1) are condition-dependent, and despite some conflicting reports on the physiological impact of *PYC1* and *PYC2* deletion (Brewster *et al*. 1994; Knijnenburg *et al*. 2009; Pronk *et al*. 1996; Stucka *et al*. 1991), one study shows that only deletion of *PYC1* leads to aspartate auxotrophy (Brewster *et al*. 1994). *PYC2* was therefore deleted. Deletion of *MAE1*, which encodes a mitochondrial malic enzyme catalyzing the oxidative decarboxylation of malate to pyruvate, does not show a clear phenotype. However, double deletion of *MAE1* and *PYK2* reduces the specific growth rate on ethanol by 62 % (Boles *et al*. 1998). As *PYK2* was deleted in the MG strain, *MAE1* was retained. The gluconeogenic enzymes PEP carboxykinase (Pck1) and fructose-1,6-bisphosphatase (Fbp1) are essential for bypassing the irreversible pyruvate-kinase and phosphofructokinase reactions, respectively, during growth on non-fermentable carbon sources (Soontorngun *et al*. 2007; Valdés-Hevia *et al*. 1989).

*CIT3, SDH1b, SHH3, SHH4 and PYC2* were deleted in two consecutive transformations rounds in the naïve reference strain and the CCMin1 strain (glyc^min^ fer^min^ ppp^min^), resulting in IMX1805 (tca^min^) and IMX1806 (CCMin2: glyc^min^ fer^min^ ppp^min^ tca^min^), respectively. Both strains grew as well as their parental strains in chemically defined medium supplemented with glucose or ethanol (Fig. 3).

#### Mitochondrial Carriers

The 35 nuclearly encoded mitochondrial carriers (MCs) mediate transport of numerous metabolites, nucleotides, cofactors and inorganic anions between mitochondrial matrix and cytosol (Palmieri *et al*. 2006). Based on extensive functional analysis studies (Palmieri and Monné 2016; Palmieri *et al*. 1996), 19 MCs involved in transport of pyruvate, TCA-cycle intermediates, CoA, ADP, ATP, Pi, NAD^+^, FAD and thiamine pyrophosphate (cofactor of pyruvate dehydrogenase and α-ketoglutarate dehydrogenase) were considered to be part of CCM (Suppl. Table S2, Fig. 2). Potential genetic redundancy was identified for 10 of these MCs, with protein sequence similarity varying between 51 and 87 % (Suppl. Table S1). In addition to genetic redundancy, functional redundancy has to be considered since several genetically distinct transporters can transport the same solutes, as exemplified by the antiport of ADP and ATP across the mitochondrial membrane by three Aac homologs as well as by Sal1. Aac2 and Aac3 originate from WGD while Aac1 does not. Sal1 shares no homology with the Aac carriers and harbours an additional Ca^2+^-binding domain (Cavero *et al*. 2005; Smith and Thorsness 2008). Several studies indicate Aac2 as major paralog, whose presence suffices to sustain adenine nucleotide transport during respiratory growth (Adrian *et al*. 1986; Cavero *et al*. 2005; Chen 2004; Gawaz *et al*. 1990; Knijnenburg *et al*. 2009; Laco *et al*. 2010; Smith and Thorsness 2008). *AAC1*, *AAC3* and *SAL1* were therefore all candidates for deletion. NAD^+^, synthesized in the cytosol and required for the NAD^+^-dependent mitochondrial dehydrogenases in CCM, is imported by two MCs encoded by *NDT1* and *NDT2*, paralogs with 51 % similarity at the protein level (Suppl. Table S1). *NDT1* and *NDT2* are individually dispensable for growth on glucose or ethanol, but deletion of both precludes growth on non-fermentable carbon sources (Todisco *et al*. 2006). Therefore, only one of the paralogs, *NDT2* was chosen for deletion. Since import of FAD, CoA and thiamine pyrophosphate is crucial for mitochondrial activity, the corresponding unique genes (*FLX1*, *LEU5* and *TPC1*) were retained (Bafunno *et al*. 2004; Marobbio *et al*. 2002; Prohl *et al*. 2001; Tzagoloff *et al*. 1996). Pyruvate is located at the interface of glycolysis and the TCA cycle and, in addition, mitochondrial pyruvate is required for synthesis of branched-chain amino acids (BCAA). Pyruvate import into mitochondria is mediated by three isoenzymes: Mpc1, Mpc2 and Mpc3. Mpc1 is constitutively expressed and forms complexes with either of the highly homologous Mpc2 or Mpc3 isoenzymes (Bender *et al*. 2015). *MPC2* is expressed during fermentative growth, while *MPC3* is expressed during respiratory growth. Deletion of *MPC2* leads to a severe growth defect, even in glucose-containing medium supplied with BCAA, while *MPC3* deletion leads to a modest (20 %) decrease of specific growth rates on non-fermentable carbon sources (Herzig *et al*. 2012; Timón-Gómez *et al*. 2013). Based on these literature data, it was decided to delete *MPC3*.

Sfc1 and Dic1 employ different mechanisms to import succinate into mitochondria and, since both are essential for growth on ethanol (Fernández *et al*. 1994; Palmieri *et al*. 1997; Palmieri *et al*. 1999b), neither were eliminated. Oxaloacetate is mainly transported by Oac1, whose removal only has a minor impact on specific growth rate on glucose medium, which is linked to its secondary function as an exporter of α-isopropylmalate for leucine biosynthesis (Marobbio *et al*. 2008; Palmieri *et al*. 1999a). Since we observed a 26 % reduction of the specific growth rate on glucose upon deletion of *OAC1* in the CEN.PK genetic background used in this study (Suppl. Fig. S1), it was retained in strain construction.

Four additional and partially functionally redundant MCs with different transport mechanisms and affinities mediate organic acid transport. Ctp1 is a citrate/malate antiporter, the paralogous carriers Odc1 and Odc2 are α-ketoglutarate/oxodicarboxylate antiporters and Yhm2 exchanges α-ketoglutarate and citrate, thereby enabling NADPH shuttling between cytosol and mitochondria (involving isocitrate dehydrogenase and aconitase) (Castegna *et al*. 2010; Cho *et al*. 1998; Kaplan *et al*. 1995; Kaplan *et al*. 1996; Palmieri *et al*. 2001; Scarcia *et al*. 2017; Tibbetts *et al*. 2002). Deletion of *CTP1* or double deletion of *ODC1* and *ODC2* does not affect growth, while triple deletion of *YHM2, ODC1* and *ODC2* does (Scarcia *et al*. 2017; Tibbetts *et al*. 2002). Based on these literature data, *CTP1, ODC1* and *ODC2* were selected for deletion, with the realization that their combined deletion might affect di- and tri-carboxylic acids trafficking.

In total, 8 MCs were targeted for elimination. First, *AAC1*, *AAC3*, *SAL1* and *MPC3* were simultaneously deleted, followed by simultaneous deletion of *NDT2*, *CTP1, ODC1* and *ODC2*. Deletion of *AAC1*, *AAC3*, *SAL1* and *MPC3* in the naïve reference strain, resulting in strain IMX2360 only marginally affected specific growth rate on SMD (3-5 % decrease), but had a stronger impact on growth on SME (14-18 % slower growth, Fig 3). These results are in agreement with the reported roles of these MCs in respiratory growth. When introduced in CCMin2 (glyc^min^ fer^min^ ppp^min^ tca^min^), resulting in strain IMX1984, the same set of deletions did not affect specific growth rate on either SMD or SME.

Combined deletion of *NDT2*, *CTP1, ODC1* and *ODC2* reduced specific growth rate on SMD by 23 % in the naïve reference strain (resulting in strain IMX2230) and 30 % in IMX1984 (glyc^min^ fer^min^ ppp^min^ tca^min^ *aac1Δ aac3Δ sal1Δ mpc3Δ*; resulting in strain IMX2231) (Fig 3). When, instead, only *NDT2, CTP1* and *ODC1* were deleted in the naïve reference strain (resulting in strain IMX2404) or in engineered background strain IMX1984 (resulting in strain IMX2407, called CCMin3: glyc^min^ fer^min^ ppp^min^ tca^min^ mc^min^), specific growth rate on SMD was not affected and only a small (3-7 %) reduction of growth rate was observed on SME (Suppl. Fig S2 and Fig 3). CCMin3 (glyc^min^ fer^min^ ppp^min^ tca^min^ mc^min^), retained 96 % of the specific growth rate of the reference strain on SMD and 82 % of its specific growth rate on SME.

#### Fumarate reductases, acetyl-CoA synthesis and glyoxylate cycle

Cytosolic (Frds1) and mitochondrial (Osm1) fumarate reductases re-oxidise FADH2 which has been proposed to be important for protein folding under anaerobic conditions (Camarasa *et al*. 2007; Jouhten and Penttilä 2014; Liu *et al*. 2013; Neal *et al*. 2017). Double deletion of *FRDS1* and *OSM1* has no phenotypic effect on complex glucose medium under aerobic conditions (Arikawa *et al*. 1998). However, Osm1 has a moonlighting function outside CCM, as it contains two translation sites, leading to the targeting to the ER of an Osm1 variant. Therefore, only *FRDS1* was considered for deletion in the design of a minimal CCM strain.

The glyoxylate cycle, which is essential for providing biosynthetic precursors with more than 2 carbon atoms during growth on fatty acids and two-carbon compounds, encompasses reactions in the peroxisome and cytosol (Kunze *et al*. 2006; Xiberras *et al*. 2019) and uses acetyl-CoA as substrate made by the acetyl-CoA synthesis pathway. Ethanol is converted into to acetyl-CoA via alcohol dehydrogenase (already reduced in the MG strain), acetaldehyde dehydrogenases and acetyl-CoA synthetases. Five acetaldehyde dehydrogenase isoenzymes, Ald2 to Ald6, oxidize acetaldehyde to acetate with either NADP^+^ or NAD^+^ as cofactor. The mitochondrial isoenzymes Ald4 and Ald5, required for growth on ethanol (Boubekeur *et al*. 2001; Kurita and Nishida 1999) and for maintenance of a functional respiratory chain (Kurita and Nishida 1999), were both retained. Ald6 is the major cytosolic isoenzyme, whose elimination strongly affects growth on fermentable and non-fermentable carbon sources (Meaden *et al*. 1997). The other two cytosolic acetaldehyde dehydrogenases, Ald2 and Ald3 are involved in conversion of 3-aminopropanal to β-alanine for pantothenic acid biosynthesis (Meaden *et al*. 1997; White *et al*. 2003). As single deletion of *ALD2* or *ALD3* does affect growth on ethanol or glucose and *ALD2* is the major paralog in pantothenic acid production (Navarro-Aviño *et al*. 1999; White *et al*. 2003), *ALD3* was considered for deletion.

Acetate is then converted to acetyl-CoA via Acs1, whose localisation is under debate. Acs1 having been reported to occur in the cytosol, the nucleus and in peroxisomes, depending on growth conditions (Chen *et al*. 2012; Krivoruchko *et al*. 2015). Acs1 and its isoenzyme Acs2 are essential for growth on non-fermentable and fermentable carbon sources, respectively (Chen *et al*. 2012). The mitochondrial acetyl-CoA hydrolase Ach1, is also able to convert acetate into acetyl-CoA, but uses succinyl-CoA as CoA donor. Deletion of *ACH1* leads to reduced chronological lifespan, severe mitochondrial damage and accumulation of reactive oxygen species (Orlandi *et al*. 2012; Takahashi *et al*. 2006). *ACS1*, *ACS2* and *ACH1* were therefore retained.

The glyoxylate cycle is initiated by Cit2, an extramitochondrial isoenzyme of the mitochondrial Cit1 and Cit3 citrate synthases, whose localisation is under debate and has been reported in the cytosol and peroxisome (Chen *et al*. 2012; van Rossum *et al*. 2016a). Citrate is then converted into isocitrate in the cytosol by the dually localized enzyme Aco1 in cytosol and mitochondria (Regev-Rudzki *et al*. 2005). Via a series of cytosolic and peroxisomal reactions (some localisations under debate), including: the isocitrate lyase Icl1 (cytosol) (Chaves *et al*. 1997; Taylor *et al*. 1996), the malate synthase (Mls1/Mls2 cytosol and peroxisome) (Chen *et al*. 2012; Hiltunen *et al*. 2003; Kunze *et al*. 2002) and malate dehydrogenase (Mdh2 in cytosol and Mdh3 occurs in the peroxisome) (Hiltunen *et al*. 2003), the net synthesis of TCA-cycle intermediates is enabled from acetyl-CoA.

Possible redundancies of glyoxylate enzymes also involved in TCA cycle were discussed above, with only Cit3 selected for elimination in a minimal CCM strain. Since the glyoxylate cycle enzymes Cit2, Mls1, Icl1 and Mdh2 (cytosolic isoenzyme of the mitochondrial Mdh1) are either essential for growth on C_2_-compounds or their elimination leads to strong reductions in growth rate, they were retained in the minimal CCM design (Chen *et al*. 2012; Fernández *et al*. 1992; Hartig *et al*. 1992; Kim *et al*. 1986; Kunze *et al*. 2006; Lee *et al*. 2011; Minard and McAlister-Henn 1991). The proteins Icl2, Mls2 and Mdh3 are homologous to Icl1, Mls1 and Mdh1/Mdh2, respectively, but have (additional) functions outside CCM (Fernández *et al*. 1993; Hartig *et al*. 1992; Luttik *et al*. 2000) and were therefore also retained.

The peroxisomes harbour the NADP^+^-dependent isocitrate dehydrogenase Idp3. Deletion of its mitochondrial homologs Idp1 and Idp2 does not affect growth on ethanol or glucose (Haselbeck *et al*. 1992; Zhao and McAlister-Henn 1996). Idp1 and Idp2 were therefore the only genes considered for elimination in the minimal CCM design.

Triple deletion of *FRDS1, IDP1* and *IDP2* in the naïve reference strain (resulting in strain IMX2470: fum^min^ glyox^min^) did not affect specific growth rate on SMD and caused a 7 % lower growth rate on SME (Fig.3 and Suppl. Fig. S2). Deletion in CCMin3 (IMX2407: glyc^min^ fer^min^ ppp^min^ tca^min^ mc^min^) did not affect specific growth rate on either SMD or SME (Fig. 3). The resulting strain CCMin4 (IMX2475: glyc^min^ fer^min^ ppp^min^ tca^min^ mc^min^ fum^min^ glyox^min^) retained 94 % and 79 % of the specific growth rate of the naïve reference strain IMX581, on SMD and SME, respectively. For reasons of experimental efficiency, *ALD3* was removed in the final deletion round (see below).

#### Glycerol synthesis

Glycerol production is essential for redox balancing in anaerobic *S. cerevisiae* cultures (Van Dijken and Scheffers 1986). In addition, glycerol plays a key role in osmotolerance and maintenance of cellular volume and turgor pressure during growth under hypertonic conditions (Ansell *et al*. 1997; Nevoigt and Stahl 1997). The conversion of dihydroxyacetone phosphate to glycerol-3-phosphate is catalysed by the isoenzymes Gpd1 and Gpd2. *gpd1* deletion mutants are osmosensitive, but show no growth defects in the absence of stress (Al-Saryi *et al*. 2017; Albertyn *et al*. 1994; Björkqvist *et al*. 1997). In contrast, *gpd2* null mutants show a mild reduction of aerobic growth rates and strongly decreased growth rates under anaerobic conditions (Eriksson *et al*. 1995; Nissen *et al*. 2000). Glycerol-3-phosphate is converted into glycerol by the redundant Gpp1 and Gpp2 isoenzymes. Single deletion of either enzyme neither affects osmotolerance nor growth on glucose or ethanol, while *gpp1* mutants have been reported to show extended lag phases in anaerobic cultures (Norbeck *et al*. 1996; Pahlman *et al*. 2001). Therefore, *GPD1* and *GPP2* were chosen for deletion.

Triple deletion of *ALD3, GPD1* and *GPP2* did not significantly affect specific growth rate on SMD, while a small growth rate reduction was observed on SME in both the naïve reference strain and the engineered background (IMX2509: Ace^,min^ Glycerol^min^ and IMX2519/CCMin5: glyc^min^ fer^min^ ppp^min^ tca^min^ mc^min^ fum^min^ glyox^min^ Ace^,min^ Glycerol^min^, respectively Fig. 3). The lower specific growth rate on SME could be attributed to the double deletion of *GPD1* and *GPP2* (strain IMX2612, Fig. 3 and Suppl. Fig. S2).

The auxotrophic 35-deletion strain IMX2519 (glyc^min^ fer^min^ ppp^min^ tca^min^ mc^min^ fum^min^ glyox^min^ Ace^,min^ Glycerol^min^) grew at 93 % and 71 % of the specific growth rate of control strain IMX581 on uracil-supplemented SMD and SME, respectively. Integration of a *URA3* cassette yielded the uracil-prototrophic 35-deletion strain IMX2538, which was labelled ‘minimal CCM strain’. This prototrophic strain grew at 94 % of the prototrophic control strain with a full set of CCM genes CEN.PK113-7D on SMD and at 76 % on SME (Fig. 3). These values were within the 25 % boundary that were initially set, and the physiology of the minimal CCM strain was further explored.

### A *S. cerevisiae* strain with minimalized CCM shows only mild growth defects on synthetic media

The genome sequence of the minimal CCM strain was analysed by short-read and long-read techniques. Long-read sequencing revealed that 9 transformation rounds and deletion of 22 genes from the MG strain had not led to chromosomal rearrangements or deletions. Previously reported duplicated regions on chromosomes 3 and 5 of the MG strain based on karyotyping and short read sequencing (Solis-Escalante *et al*. 2015) were also observed in this study with long-read sequencing. Sequence analysis confirmed that all 22 targeted CCM genes were correctly deleted from the MG strain. The genome of the minimal CCM strain showed 45 Single Nucleotide Polymorphisms (SNPs) relative to the MG strain, of which eight were located in genes and only four led to an amino acid change (Table 1), none of which affected proteins involved in CCM.

**Table 1.** Single-nucleotide mutations identified in coding regions of the prototrophic minimal CCM strain IMX2538. Single-nucleotide changes in *S. cerevisiae* IMX2538 (prototrophic minimal CCM) relative to the genome sequence of *S. cerevisiae* IMX372 (prototrophic minimal glycolysis (MG)) (Solis-Escalante *et al*. 2015).

| Systematic name | Name | Type* | Amino acid change |
| --- | --- | --- | --- |
| YBR114W | *RAD16* | NS | Ile-202-Thr |
| YDL035C | *GPR1* | S | Asn-523-Asn |
| YDR098C | *GRX3* | NS | Glu-239-Asp |
| YFL062W | *COS4* | S | Cys-151-Cys |
| YLR002C | *NOC3* | NS | Asp-526-Glu |
| YML058W | *SML1* | NS | Gly-52-Ser |
| YMR154C | *RIM13* | S | Lys-265-Lys |
| YNL273W | *TOF1* | S | Asn-117-Asn |

\*S: synonymous, NS: non-synonymous

The physiology of the minimal CCM strain was compared to that of the congenic reference strain CEN.PK113-7D, which has a full complement of CCM genes, in pH-controlled aerobic bioreactor cultures on SMD. Consistent with the analyses in shake flasks, the specific growth rate of the minimal CCM strain in these cultures was 8 % lower than the level of the reference strain CEN.PK113-7D (Table 2). During the glucose consumption phase, biomass-specific glucose and oxygen consumption rates of the two strains, as well as their ethanol and CO_2_ production rates and their biomass and ethanol yields on glucose were also similar. The minimal CCM strain did exhibit a higher acetate production rate and yield (63 % and 71 % higher, respectively) than the reference strain, a difference already observed for the MG strain (Solis-Escalante *et al*. 2015). Similarly, a lower glycerol production rate and glycerol yield on glucose (27 % and 23 % lower, respectively), was in line with data reported for a *gpd1* deletion mutant (Björkqvist *et al*. 1997). After the diauxic shift, growth of the minimal CCM strain on ethanol, glycerol and organic acids produced during the glucose phase, proceeded at a 17 % lower rate than observed for the reference strain (Table 2). As a macroscopic characterization based on extracellular products might mask subtle differences of intracellular fluxes, intracellular concentrations of CCM intermediates were measured during the mid-exponential growth phase on glucose. The concentrations of these metabolites hardly differed between the minimal CCM and the control strains (Table 3). These results indicated that a 32 % reduction of the complement of genes encoding CCM enzymes of *S. cerevisiae* had only a small impact on its physiology under standard laboratory conditions.

**Table 2.** Physiological characterization of a 35-deletion, minimal CCM prototrophic *S. cerevisiae* strain in aerobic bioreactor batch cultures. *S. cerevisiae* strains were grown at pH 5.0 and at 30 °C in aerobic bioreactors on synthetic medium with glucose as sole carbon source. Data are presented as average and standard deviation of 3 biological replicates for *S. cerevisiae* strains CEN.PK113-7D (naïve reference) and IMX2538 (minimal CCM). Data for S. cerevisiae IMX372 (minimal glycolysis) were recalculated from the raw data of Solis-Escalante *et al*. (2015) obtained with two biological replicates. Statistical significance with respect to CEN.PK113-7D is represented in bold with an asterisk (two-tailed t-test, equal variances, P<0.05).

|  | CEN.PK113-7D<br>(Naïve reference) | IMX372<br>(Minimal Glycolysis) | IMX2538<br>(Minimal CCM) |
| --- | --- | --- | --- |
| **Glucose phase** |  |  |  |
| $\mu_{\max}$ (h<sup>-1</sup>) | 0.37 ± 0.00 | **0.38 ± 0.01\*** | **0.34 ± 0.00\*** |
| $q_s$ (mmol.g<sub>DW</sub><sup>-1</sup>.h<sup>-1</sup>) | -16.2 ± 0.2 | -15.7 ± 0.7 | -15.4 ± 0.5 |
| $q_{\text{Ethanol}}$ (mmol.g<sub>DW</sub><sup>-1</sup>.h<sup>-1</sup>) | 23.5 ± 1.5 | 23.1 ± 1.1 | 23.2 ± 2.1 |
| $q_{\text{Glycerol}}$ (mmol.g<sub>DW</sub><sup>-1</sup>.h<sup>-1</sup>) | 1.52 ± 0.05 | 1.40 ± 0.02 | **1.11 ± 0.05\*** |
| $q_{\text{Acetate}}$ (mmol.g<sub>DW</sub><sup>-1</sup>.h<sup>-1</sup>) | 0.44 ± 0.03 | **0.82 ± 0.04\*** | **0.71 ± 0.02\*** |
| $q_{\text{CO}_2}$ (mmol.g<sub>DW</sub><sup>-1</sup>.h<sup>-1</sup>) | 23.4 ± 0.2 | | 22.6 ± 0.5 |
| $q_{\text{O}_2}$ (mmol.g<sub>DW</sub><sup>-1</sup>.h<sup>-1</sup>) | -6.8 ± 0.4 | | -7.0 ± 0.2 |
| $Y_{\text{biomass/glucose}}$ (g<sub>DW</sub>·g<sub>glucose</sub><sup>-1</sup>) | 0.13 ± 0.00 | 0.13 ± 0.00 | 0.12 ± 0.00 |
| $Y_{\text{ethanol/glucose}}$ (mol.mol<sup>-1</sup>) | 1.45 ± 0.09 | 1.48 ± 0.01 | 1.51 ± 0.09 |
| $Y_{\text{glycerol/glucose}}$ (mol.mol<sup>-1</sup>) | 0.09 ± 0.00 | 0.09 ± 0.01 | **0.07 ± 0.00\*** |
| $Y_{\text{acetate/glucose}}$ (mol.mol<sup>-1</sup>) | 0.03 ± 0.00 | 0.05 ± 0.00 | **0.05 ± 0.00\*** |
| **Post-diauxic phase** |  |  |  |
| $\mu_{\max}$ (h<sup>-1</sup>) | 0.10 ± 0.00 | **0.12 ± 0.00\*** | **0.08 ± 0.01\*** |
| $q_{\text{Ethanol}}$ (mmol.g<sub>DW</sub><sup>-1</sup>.h<sup>-1</sup>) | -3.10 ± 0.19 | **-3.93 ± 0.04\*** | -3.07 ± 0.31 |

**Table 3.** Intracellular metabolite profiles of a 35-deletion, minimal CCM prototrophic *S. cerevisiae* strain in aerobic bioreactor batch cultures. Intracellular metabolite contents were measured during the mid-exponential glucose phase of aerobic bioreactor batch cultures of *S. cerevisiae* CEN.PK113-7D (naïve reference) and IMX2538 (minimal CCM) (see Table 2 for other physiological data). Data represent average and standard deviation of data from analyses on three independent cultures for each strain. Fold differences that are statistically significant are indicated in bold with an asterisk (two-tailed t-test, equal variances, P<0.05). # below detection limit.

| Metabolite | CEN.PK113-7D<br>(naïve reference) | IMX2538<br>(Minimal CCM) | Fold<br>difference |
| --- | --- | --- | --- |
| $\mu\text{mol.}(\text{g biomass dry weight})^{-1}$ | | | |
| **Glycolysis** |  |  |  |
| Glucose 6-phosphate | $4.13 \pm 0.49$ | $5.01 \pm 0.38$ | 1.2 |
| Fructose 6-phosphate | $0.38 \pm 0.06$ | $0.59 \pm 0.08$ | **1.5\*** |
| Fructose 1,6-bisphosphate | $17.53 \pm 0.82$ | $20.52 \pm 0.66$ | **1.2\*** |
| Glyceraldehyde 3-phosphate | $0.15 \pm 0.01$ | $0.19 \pm 0.01$ | **1.3\*** |
| Dihydroxyacetone phosphate | $0.79 \pm 0.13$ | $2.08 \pm 0.66$ | 2.6 |
| 3-Phosphoglycerate | $1.30 \pm 0.12$ | $2.52 \pm 0.28$ | **1.9\*** |
| 2-Phosphoglycerate | $0.16 \pm 0.01$ | $0.17 \pm 0.01$ | 1.0 |
| Phosphoenolpyruvate | $0.25 \pm 0.01$ | $0.12 \pm 0.01$ | **0.5\*** |
| **Trehalose synthesis** |  |  |  |
| Trehalose 6-phosphate | $9.34 \pm 1.18$ | $14.48 \pm 1.67$ | **1.6\*** |
| Trehalose<sup>#</sup> | $0.20 \pm 0.04$ | $0.41 \pm 0.09$ | **2.1\*** |
| **Pentose phosphate pathway** |  |  |  |
| 6-Phosphoglucononate | $1.01 \pm 0.02$ | $1.11 \pm 0.07$ | 1.1 |
| Ribose 5-phosphate | $0.40 \pm 0.02$ | $0.51 \pm 0.03$ | **1.3\*** |
| Ribulose 5-phosphate | $0.20 \pm 0.03$ | $0.33 \pm 0.06$ | **1.6\*** |
| Xylulose 5-phosphate | $0.43 \pm 0.07$ | $0.72 \pm 0.13$ | 1.7 |
| Sedoheptulose 7-phosphate | $0.50 \pm 0.06$ | $0.54 \pm 0.04$ | 1.1 |
| Erythrose 4-phosphate | $0.004 \pm 0.000$ | $0.005 \pm 0.000$ | **1.3\*** |
| **TCA cycle and Glyoxylate cycle** |  |  |  |
| Citrate | $4.56 \pm 0.65$ | $5.80 \pm 0.98$ | 1.3 |
| Isocitrate | $0.02 \pm 0.00$ | $0.03 \pm 0.00$ | **1.7\*** |
| α-ketoglutarate | $0.29 \pm 0.02$ | $0.67 \pm 0.24$ | 2.3 |
| Succinate | $0.37 \pm 0.03$ | $0.37 \pm 0.12$ | 1.0 |
| Fumarate | $0.10 \pm 0.02$ | $0.18 \pm 0.06$ | 1.8 |
| Malate | $0.60 \pm 0.08$ | $1.03 \pm 0.19$ | **1.7\*** |
| **Nucleotides and cofactors** |  |  |  |
| Energy charge | $0.86 \pm 0.00$ | $0.88 \pm 0.01$ | 1.0 |
| AMP | $0.13 \pm 0.01$ | $0.17 \pm 0.01$ | **1.3\*** |
| ADP | $1.55 \pm 0.08$ | $1.43 \pm 0.11$ | 0.9 |
| ATP | $5.50 \pm 0.16$ | $5.58 \pm 0.21$ | 1.0 |
| UDP | 0.23 ± 0.01 | 0.22 ± 0.05 | 0.9 |
| UTP | 1.61 ± 0.06 | 1.57 ± 0.10 | 1.0 |
| GDP | 0.27 ± 0.02 | 0.21 ± 0.07 | 0.8 |
| GTP | 1.58 ± 0.06 | 1.49 ± 0.15 | 0.9 |
| CDP | 0.13 ± 0.02 | 0.15 ± 0.01 | 1.1 |
| CTP | 0.84 ± 0.02 | 0.82 ± 0.03 | 1.0 |
| NADH | 0.18 ± 0.02 | 0.15 ± 0.02 | 0.8 |
| NAD<sup>+</sup> | 85.43 ± 1.90 | 91.82 ± 2.64 | **1.1\*** |
| NADP<sup>+</sup> | 8.49 ± 0.67 | 6.87 ± 0.25 | **0.8\*** |
| Acetyl-CoA | 5.80 ± 0.18 | 5.38 ± 0.34 | 0.9 |
| FAD | 0.87 ± 0.12 | 0.54 ± 0.18 | 0.6 |

### Dissecting individual from synergistic responses to growth on a range of conditions

To explore genetic redundancy of CCM genes, the minimal CCM strain and the congenic reference strain CEN.PK113-7D were grown under a broad range of conditions. Some of these were chosen based on previously reported phenotypes (e.g. high osmolarity) or connection to CCM (e.g. growth on various carbon sources), while others subjected the strains to adverse conditions (e.g. acidic or alkaline pH).

Consistent with reports that deletion of *GPD1* causes decreased osmotolerance (Albertyn *et al*. 1994; Eriksson *et al*. 1995), the minimal CCM strain grew 13 to 25% slower than the reference strain exposed to high osmolarity, which was imposed by adding high concentrations of sorbitol (1M and 2M) or glucose (10 % and 20 % w/v). Construction and analysis of strains with different combinations of deletions in reference and CCM-minimization backgrounds confirmed that this growth reduction specifically resulted from *GPD1* deletion, rather than from synergistic effects (Fig. 4 and Suppl. Fig. S3). *MPC3* deletion has been reported to cause lower growth rates on glycerol or lactate as sole carbon source (Timón-Gómez *et al*. 2013). Since Mpc3 is a pyruvate transporter, growth was assessed directly on chemically defined medium with pyruvate as sole carbon source. The minimal CCM strain grew 79% slower than the control strain, however this could surprisingly not be attributed to the *MPC3* deletion (Fig. 4 and Suppl. Fig. S3).

**Figure 4.**
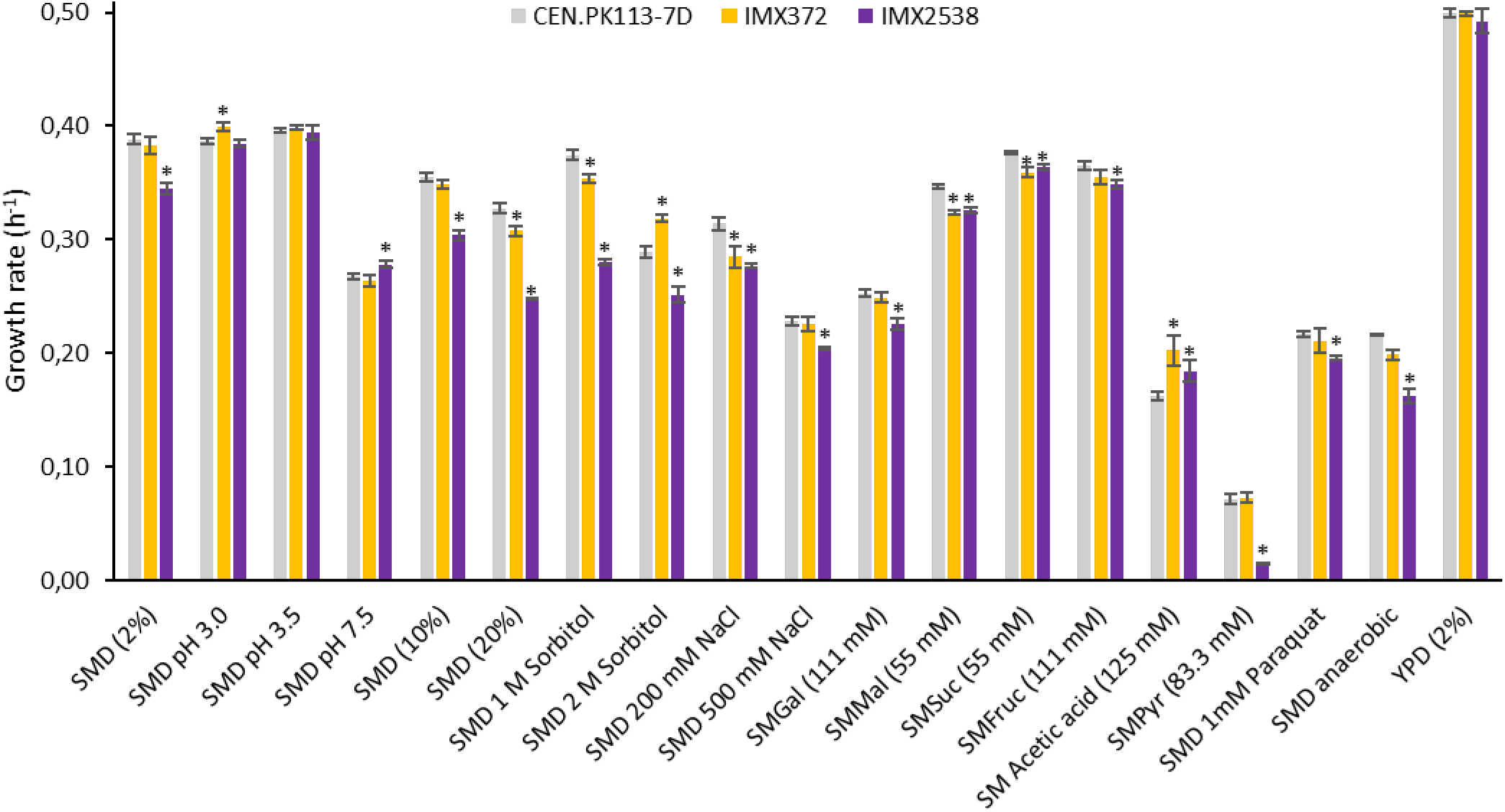
Specific growth rates of 35-deletion, prototrophic minimal CCM strain under a broad range of growth conditions. Specific growth rates of the prototrophic *S. cerevisiae* strains CEN.PK113-7D (naïve reference strain), IMX372 (minimal glycolysis (MG)) and IMX2538 (minimal CCM) under different growth conditions. Specific growth rates were measured in triplicate cultures using a growth profiler, except for those in SMPyr and SMD-anaerobic, which were measured in independent duplicate shake flask cultures. Abbreviations indicate the following growth conditions: SM, synthetic medium; SMD, synthetic medium with glucose; Gal, galactose as carbon source; Mal, maltose as carbon source; Suc, sucrose as carbon source; Fruc, fructose as carbon source; Pyr, pyruvate as carbon source; YPD, complex medium with glucose. Significant changes in growth rate of IMX372 and IMX2538 with respect to CEN.PK113-7D are indicated with a * (two-tailed paired homoscedastic t-test p<0.05).

Combined deletion of *AAC1*, *AAC3*, *MPC3* and *SAL1* caused a 3 – 5 % and a 14 – 18 % decrease of growth rates on SMD and SME, respectively, thus marking their importance on these carbon sources. According to previous reports, individual deletion of these four genes does not affect growth on glucose and individual deletion of *AAC1*, *AAC3* and *SAL1* does not affect growth on ethanol (Cavero *et al*. 2005; Kolarov *et al*. 1990; Smith and Thorsness 2008; Timón-Gómez *et al*. 2013). Deletion of *MPC3* has been reported to cause a decrease in growth rate on glycerol and lactate (Timón-Gómez *et al*. 2013), and may therefore also be responsible for the lower growth rate on ethanol. Reintroduction of *MPC3* in strain IMX1984 (glyc^min^ fer^min^ ppp^min^ tca^min^ *aac1Δ aac3Δ sal1Δ mpc3Δ*) increased specific growth rate on SME by only 3 %, while reintroduction in IMX2519 (CCMin 5: glyc^min^ fer^min^ ppp^min^ tca^min^ mc^min^fum^min^ glyox^min^ Ace^,min^ Glycerol^min^) did not affect growth rate (Suppl. Fig. S3). These results suggest that the observed impact of quadruple deletion of *AAC1*, *AAC3*, *MPC3* and *SAL1* on ethanol growth was caused by synergistic effects.

Some paralogs have been reported to be specifically important under anaerobic conditions (*AAC3, FRDS1*) (Arikawa *et al*. 1998; Kolarov *et al*. 1990). In line with these reports, while the MG strain showed the same growth rate as CEN.PK113-7D under anaerobic conditions, the minimal CCM strain showed a 25 % lower anaerobic growth rate (Fig. 4). However, testing of deletions in the reference background indicated that this difference was not caused by the deletions of *AAC3* or *FRDS1* (Suppl. Fig. S3).

Over the broad range of conditions tested, including several stress conditions, few differences in specific growth rate were observed (Fig. 4). Combining the phenotype of strains with individual and clustered deletions enabled to identify synergistic interactions between minor paralogs.

## Discussion

Genetic reduction has been applied in several microorganisms (Lara and Gosset 2019), including *B. subtilis* (Morimoto *et al*. 2008), *E.coli* (Hashimoto *et al*. 2005), *Lactococcus lactis* (Zhu *et al*. 2017)*, C. glutamicum* (Suzuki *et al*. 2005), *Streptomyces* species (Komatsu *et al*. 2010), *Pseudomonas* species (Lieder *et al*. 2015) and *Schizosaccharomyces pomb*e (Sasaki *et al*. 2013), with the purpose of discovering a minimal genome content and/or for engineering efficient cell factories. In *S. cerevisiae*, Marakami *et al*. reduced genome content by 5 % by deleting 15 terminal chromosomal regions (Murakami *et al*. 2007). Moreover, the creation of a synthetic yeast genome in the Sc2.0 project is accompanied by an 8 % genome reduction by deletion of long terminal repeats, retrotransposons and introns; and engineering of a single chromosome yeast strain was characterised by a 9 % decrease in DNA (Richardson *et al*. 2017; Shao *et al*. 2018). Genome reduction studies typically target two types of DNA sequences: non-expressed DNA (cryptic genes, mobile DNA) and irrelevant/non-essential genes. These DNA elements can be targeted by random strategies for which little knowledge is required, such as transposon mutagenesis or the elegant SCRaMbLE technique used for the recent reduction of left synthetic chromosome arm XII in *S. cerevisiae* (Luo *et al*. 2021). In the present study, knowledge-based reduction of the gene complement for CCM in *S. cerevisiae* was informed by gene expression data and previous phenotypic analysis on single-knockout mutants (Luo *et al*. 2021; Murakami *et al*. 2007; Richardson *et al*. 2017; Shao *et al*. 2018).

In this study, we reduced genetic complexity of central carbon metabolism (CCM) in *S. cerevisiae* by deletion of 35 genes encoding enzymes and transporters. This reduction corresponded to elimination of 32 % of the (iso)enzymes and transporters involved in the included processes, without major impacts on strain physiology, which was tested under a broad range of conditions (Fig. 2-4). The present study built on earlier work by Solis-Escalante *et al*. (2015), who eliminated 50 % of isoenzymes involved in glycolysis and ethanolic fermentation with a similar small impact on physiology. The attainable reduction of gene sets for enzymes and transporters involved in other CCM pathways differed, with 50 % for fumarate reduction and glycerol synthesis, 37 % for the mitochondrial carriers, 36 % for the pentose-phosphate pathway, 23 % for the TCA cycle, 14% for the glyoxylate cycle, 20 % for the glycolysis – TCA cycle interface plus gluconeogenesis, and 8 % for acetyl-CoA metabolism. The lower attainable genetic reduction of the four latter pathways can be largely attributed to neofunctionalization and relocalization of enzymes during evolution.

Our results show that yeast CCM is remarkably robust to genetic reduction, in particular during growth on glucose, yeast’s favourite carbon source, but also when challenged by a broad range of growth conditions. Notable exceptions were growth on pyruvate (79 % growth rate reduction), anaerobic growth on glucose (25 % reduction), growth on ethanol (24 % reduction) and growth at high osmolarity (between 13 % and 25 % lower specific growth rates). Growth-rate reductions on ethanol and at high osmolarity could be attributed to specific genes or gene combinations, while for growth on pyruvate and anaerobic growth some hypothetical targets could be excluded. The physiological role of most deleted paralogs remains elusive. Such a situation is exemplified by *TKL2* and *NQM1*, which are paralogs of the major PPP genes *TKL1* and *TAL1*, respectively. In *S. cerevisiae* strains engineered for L-arabinose utilisation, their deletion was shown to lead to lower growth rates on this pentose sugar (Matsushika *et al*. 2012; Wisselink *et al*. 2010). Clearly, as pentoses are not natural carbon sources for *S. cerevisiae*, this role cannot have provided an evolutionary driving force for fixation of these paralogs in its genome, but it does indicate potential contribution to fitness under other, as yet unidentified growth conditions. Testing the minimal CCM strain under an even wider variety of environmental conditions, including dynamics in nutrient availability and other environmental parameters may reveal physiological roles of these and other paralogs. Alternatively, the mechanisms that fixed some paralogs during evolution may have been disrupted by relatively recent mutations or gene loss (Shen *et al*. 2018). Following this reasoning, absence of a clear phenotype of knock-out mutants may have captured a stage in the evolutionary trajectory of *S. cerevisiae* that will eventually lead to loss of a paralog, evolution towards complete sub-functionalisation, or retention of functional overlap with asymmetric divergence (Kuzmin *et al*. 2020).

In this first step towards the genetic minimization of CCM in yeast, choices had to be made on which pathways and genes were considered as part of the CCM and on criteria for redundancy. For instance, transport of NAD^+^, FAD^+^, ADP/ATP and Pi across the mitochondrial membrane was considered, while transport of NAD(P)H, which requires more complex shuttle systems (Bakker *et al*. 2001; Miyagi *et al*. 2009) was not yet included. In addition, as *S. cerevisiae* cannot synthetize carnitine (Swiegers *et al*. 2001), the carnitine shuttle system transporting acetyl-CoA across compartments was not considered. Since *CRC1*, *CAT1*, *YAT1* and *YAT2* involved in this shuttle are dispensable for growth in the absence of carnitine (Swiegers *et al*. 2001), they can be considered for further genetic reduction of the CCM. Some genes required for anaerobic growth, such as *ADH3* (Bakker *et al*. 2000; Bakker *et al*. 2001) were also retained, but could be removed if fast anaerobic growth is excluded as a criterion. Several other processes and pathways are of particular interest for development of strain platforms for modular engineering yeast CCM. In this context, glucose uptake, which involves a set of 20 hexose transporters (Wieczorke *et al*. 1999) provides and interesting target for future experiments, whose minimization can benefit from a recently constructed Hxt^0^ CRISPR kit (Wijsman *et al*. 2019). Another logical target for minimization is uptake and assimilation of (alternative) carbon sources and especially of maltose, whose metabolism is enabled by highly redundant subtelomeric genes (Brickwedde *et al*. 2018; Brown *et al*. 2010).

Genetic reduction presents a first, indispensable step towards the construction of modular yeast strains for extensive remodelling of CCM. Current demands for economically competitive cell factories, with optimized titer, rate and yield (TRY), requires extensive remodelling of the CCM for the supply of precursors, (redox) cofactors and energy-rich molecules (Aslan *et al*. 2017; François *et al*. 2020; Papagianni 2012). For instance, the extensive remodelling of the native Entner-Doudoroff (ED) glycolytic pathway into the Embden-Meyerhof-Parnas (EMP) pathway improved carotenoid synthesis in *Pseudomonas putida* (Sánchez-Pascuala *et al*. 2019). Similarly, substantial efforts are invested in remodelling yeast CCM in *S. cerevisiae* to increase the supply of cytosolic acetyl-CoA, a precursor for a wide array of attractive biomolecules (Fig. 1) (van Rossum *et al*. 2016b). Also, production of complicated chemical structures like plant natural products in *S. cerevisiae* requires extensive remodelling of the entire central carbon metabolism (Koopman *et al*. 2012; Liu *et al*. 2019; Meadows *et al*. 2016). As demonstrated by Kuijpers *et al*. (2016), genetic reduction facilitates the colocalization of sets of genes in ‘pathway clusters’ and strongly accelerates the genetic remodelling of these pathways. With this strategy, the 12 steps of glycolysis and ethanolic fermentation were rapidly and efficiently swapped with heterologous variants and enabled the implementation of an innocuous DNA and RNA watermarking method (Boonekamp *et al*. 2020). A similar strategy can be considered for remodelling CCM, with the minimal CCM strain as starting point. As recently demonstrated, 44 transcriptional-unit sized DNA fragments can be assembled in *S. cerevisiae* into specialized, synthetic supernumerary chromosomes (Postma *et al*. 2021). Since the capacity of HR was not reached, assembly of synthetic chromosomes containing the set of 76 genes encoding the minimal CCM has now become a realistic objective. Subsequent CRISPR-Cas assisted removal of the duplicate CCM genes from their native locations could then generate powerful platforms for chromosome swapping and combinatorial CCM remodelling studies. The reduction of genetic complexity demonstrated in the present study therefore not only provides new insights in genetic redundancy of CCM, but also contributes to the eventual localization of all genes required for a minimized CCM on specialized, synthetic supernumerary chromosomes that allow for extensive, combinatorial remodelling of yeast metabolism for industrial applications.

## Materials and Methods

### Strains, media and maintenance

The *Saccharomyces cerevisiae* strains used in this study are all derived from the CEN.PK family (Entian and Kötter 2007; Nijkamp *et al*. 2012a) (Suppl. Table S3). The naïve, uracil auxotrophic and Cas9 containing strain, IMX581 (Mans *et al*. 2015) and the uracil auxotrophic MG (minimal glycolysis) strain IMX370 (Solis-Escalante *et al*. 2015) were used for deletion of genes encoding enzymes or transporters involved in central carbon metabolism (CCM). The naïve uracil prototrophic strain CEN.PK113-7D was used for physiological comparison. Complex medium used for propagation of yeast strains consisted of 10 g L^-1^ Bacto yeast extract, 20 g L^-1^ Bacto peptone and 20 g L^-1^ glucose (YPD), autoclaved at 110°C for 20 min. After transformation, yeast strains were selected in synthetic medium (SM) (Verduyn *et al*. 1992) containing 3.0 g L^-1^ KH_2_PO_4_, 0.5 g L^-1^ MgSO_4_·7H_2_O, 5.0 g L^-1^ (NH_4_)_2_SO_4_ and 1.0 mL L^-1^ trace elements autoclaved at 121°C for 20 min, whereafter 1.0 mL L^-1^ of filter sterilized vitamin solution was added. Before autoclaving media were set to pH 6 by 1 M KOH addition. SM was supplemented with 20 g L^-1^ glucose (SMD) or 2 % ethanol v/v (SME) for propagation and growth characterisation. Synthetic medium was supplemented with 150 mg L^-1^ uracil for uracil auxotrophic strains. For selection of transformants carrying the amdS selection marker (Solis-Escalante *et al*. 2013), ammonium sulfate was replaced as nitrogen source with 10 mM of acetamide. For experiments on SM in the growth profiler and under anaerobic conditions, ammonium sulfate was replaced by 2.3 g L^-1^ Urea. For both media in which ammonium sulfate was replaced, 6.6 g L^-1^ K_2_SO_4_ was added. Growth was performed in 500 mL shake flask containing 100 mL medium or in 100 mL shake flasks containing 20 mL medium at 30 °C and 200 rpm in an Innova 44 Incubator shaker (New Brunswick Scientific, Edisan, NJ). Culture on solid media were incubated for 3-5 days at 30 °C.

CEN.PK113-7D, IMX372, IMX2538 and several intermediate strains were tested in the growth profiler on a variety of liquid media , containing: SM(urea) plus 2 % glucose (SMD), SMD at pH 3.0, 3.5 or 7.5, SM plus 10 % glucose (SMD (10 %)), SM plus 20 % glucose (SMD (20 %)), SMD plus 1 or 2M sorbitol, SMD plus 200 or 500 mM NaCl, SM plus 111 mM galactose (SMGal), SM plus 55 mM maltose (SMMal), SM plus 55mM sucrose (SMSuc), SM plus 111 mM fructose (SMFruc), SM plus 125 mM acetic acid, SMD plus 1 mM paraquat and YPD (2 % glucose). Growth on SM plus 83.3 mM pyruvic acid was performed in shake flasks. For anaerobic growth in shake flasks SMD was supplemented with 0.01 g L^−1^ ergosterol and 0.42 g L^−1^ Tween 80 dissolved in ethanol (SMD (2 %) anaerobic) (Verduyn *et al*. 1992).

Plasmids were propagated in and isolated from chemically competent *Escherichia coli* XL1-Blue cells, which were cultivated in Lysogeny Broth containing 10 g L^-1^ Bacto tryptone, 5.0 g L^-1^ Bacto yeast extract and 5 g L^-1^ NaCl supplemented with 100 mg L^-1^ ampicillin (LB-amp) when required. *E. coli* was cultivated in 15 mL Greiner tubes containing 5 mL medium at 37 °C and 200 rpm in an Innova 4000 Incubator shaker (New Brunswick Scientific). Bacterial cultures on solid medium were incubated overnight at 37°C.

For solid medium 20 g L^-1^ of agar was added before autoclaving. All *S. cerevisiae* and *E.coli* strains were stored at -80°C in 1 ml aliquots containing 30 % v/v glycerol in appropriate medium.

### Molecular biology techniques

Plasmids were isolated from *E. coli* using the GenElute Plasmid Miniprep Kit (Sigma-Aldrich, St. Louis, MO) or the GeneJET Plasmid Miniprep Kit (Thermo Fisher Scientific, Waltham, MA) according to the provided protocols. DNA fragments for plasmid construction or integrative DNA fragments used in yeast transformation were amplified using Phusion High-Fidelity DNA Polymerase (Thermo Fisher Scientific) according to manufacturer’s instructions, using PAGE-purified or desalted oligonucleotides (Sigma-Aldrich) depending on the application. Purification of genomic Polymerase Chain Reaction (PCR) amplified DNA was performed with the GenElute PCR Clean-Up kit (Sigma-Aldrich) or the GeneJET PCR Purification Kit (Thermo Fisher Scientific) if no a-specific products were present. When a-specific products were present or when DNA was amplified from plasmids, the DNA was purified by separation using electrophoresis on a 1 % (w/v) agarose gel (TopVision Agarose, Thermo Fisher Scientific) in 1x Tris-acetate-EDTA (TAE) buffer (Thermo Fisher Scientific) or on a 2 % (w/v) agarose gel (TopVision Agarose, Thermo Fisher Scientific) in 1x Tris-Borate-EDTA (TBE) buffer (Thermo Fisher Scientific) with subsequent purification with the Zymoclean Gel DNA Recovery kit (Zymo Research). Chemical transformation of *E. coli* XL1-Blue was performed by thawing of competent cells on ice, addition of DNA, followed by a heat shock for 40s at 42°C. Subsequently, cells were incubated on ice for 2 minutes and plated immediately on selective LB-amp plates and grown overnight at 37°C. Transformation of *S. cerevisiae* was performed using the lithium acetate/single-stranded carrier DNA/polyethylene glycol method (Gietz and Woods 2002). Single-cell lines were obtained by three consecutive re-streaks on selective solid medium. Yeast genomic DNA was extracted according to Looke *et al*. (2011), the YeaStar Genomic DNA kit (Zymo Research, Irvine, CA) according to Protocol I supplied by the manufacturer or the QIAGEN Blood & Cell Culture Kit with 100/G or 20/G Genomic-tips (Qiagen, Hilden, Germany) following the manufacturer’s recommendations. Verification of the accurate genotype of engineered *S. cerevisiae* strains and *E. coli* plasmids was done by diagnostic PCR before strain storage at -80°C. These diagnostic PCRs were performed using desalted oligonucleotides and the DreamTaq PCR master Mix (Thermo Fisher Scientific) according to the manufacturer’s protocol.

### Plasmid and strain construction

Deletions were performed using CRISPR/Cas9. CRISPR/Cas9-based genome editing of *S. cerevisiae* was performed as described by Mans *et al*.(2015) with minor alterations. Plasmids containing a single gRNA (Suppl. Table S4) were constructed via Gibson assembly with a backbone containing the marker cassette and one insert fragment containing the gRNA and the 2 µm fragment. The backbone was amplified from a pMEL plasmid (Mans *et al*. 2015) with primers 5980 and 5792 and the insert fragment was amplified with a gRNA specific primer designed with the yeastriction tool (Mans et al. 2015) and primer 5979 (primers in Suppl. Table S5). Plasmids containing two gRNAs were constructed using one backbone fragment and two insert fragments, each containing one gRNA and one half of the 2 µm fragment. Backbones were PCR amplified from the pROS plasmids (Mans *et al*. 2015) with the double-binding primer 6005. Insert fragments were obtained with the gRNA specific primers together with either primer 5974 or primer 5975 (Primers in Suppl. Table S5). The backbone and gRNA insert fragment(s) were gel purified, *DpnI* digested (Thermo Fisher Scientific) and Gibson-assembled in a final volume of 5 µl using NEBuilder HiFi DNA assembly master mix (NEB, Ipswich, MA), according to manufacturer’s instructions. Assembled plasmids were transformed and subsequently isolated from LB-amp grown *E. coli*. Correct assembly was checked using diagnostic PCR (Suppl. Table S6).

IMK588 was constructed by integrating the *KanMX* marker at the *OAC1* locus of CEN.PK113-7D. The KanMX marker with homologous flanks to *OAC1* was amplified with primers 6358 and 6359 from pUG6 (Suppl. Table S4, S7 and S8).

In order to perform CRISPR editing in the MG strain (IMX370), *Cas9* was integrated by transforming a *Cas9* and *natNT2* DNA fragment, which can assemble by homologous recombination at the *CAN1* locus. The *Cas9* fragment (*can1* flank-*Cas9* expression cassette-SHR A) was PCR amplified with primers 2873 and 4653 from plasmid p414-*TEF1p-Cas9-CYC1t* (Suppl. Table S4 and S9). The *natNT2* fragment (SHR A-NatNT2 marker cassette-*can* 1 flank) was amplified with primers 3093 and 5542 from plasmid pUG-*natNT2* (Suppl. Table S4 and S9). The *Cas9* containing MG strain was stocked as IMX1331.

For genome editing using CRISPR, *S. cerevisiae* strains were transformed with 1 µg of each gRNA plasmid and 1 µg of each 120 bp double-stranded DNA repair fragment. These repair fragments were made by annealing of complimentary oligonucleotides listed in Suppl. Table S7, and consisted 60 bp homology sequences immediately upstream of the start codon and downstream of the stop codon of the targeted gene, unless stated otherwise. Transformants were plated on selective medium. Gene deletion was verified by diagnostic colony PCR on randomly picked colonies by using the primers which bind outside of the targeted open reading frame (Suppl. Table S8). gRNA plasmids were removed by growing the colonies in liquid YPD medium and subsequent plating on solid YPD medium. Plasmid removal was confirmed by growth on selective and non-selective solid media after which the strains were stored. For all transformations the corresponding gRNA plasmids and repair fragment are summarised in Suppl. Table S10.

To obtain a prototrophic strain with 35 deletions, a *URA3* transcriptional unit amplified from CEN.PK113-7D DNA (primers: 17752 and 17753, Suppl. Table S7) was integrated at the *GPP2* locus of strain IMX2520 (34 deletions). The flanks of the *URA3* repair fragment were homologous to the 60 bp immediately upstream and downstream of the *GPP2* ORF. The prototrophic 35 deletion strain IMX2538 was checked by diagnostic PCR, short-read sequencing and long-read nanopore sequencing. Integration of the *MPC3* transcriptional unit at the *X2* locus of IMX1984 and IMX2519 was achieved by amplifying the respective fragment from CEN.PK113-7D genomic DNA (primers:18025 and 18026 Suppl. Table S7) and integration by CRISPR/Cas9 using gRNA plasmid pUDR376, resulting in strains IMX2640 and IMX2641 respectively. Correct integration was verified by diagnostic PCR (Suppl. Table S8).

### Sequencing

High-quality genomic DNA of yeast for sequencing was extracted using the QIAGEN Blood & Cell Culture Kit with 100/G or 20/G Genomic-tips (QIAGEN) according to the manufacturer’s instructions. DNA concentration was measured using the BR ds DNA kit (Invitrogen, Carlsbad, CA) and a Qubit® 2.0 Fluorometer (Thermo Fisher Scientific). The purity was verified with a Nanodrop 2000 UV-Vis Spectrophotometer (Thermo Fisher Scientific).

### Short read sequencing

IMX2538 (35 deletions prototrophic strain) was sequenced using 300 bp paired-end sequencing reads prepared with the MiSeq Reagent Kit v3 on an Illumina MiSeq sequencer (Illumina, San Diego, CA). To this end, extracted DNA was mechanically sheared to 550 bp with the M220 ultrasonicator (Covaris, Wolburn, MA) and subsequently, the TruSeq DNA PCR-Free Library Preparation kit (Illumina) was employed to make a six-strain library. The samples were quantified by qPCR on a Rotor-Gene Q PCR cycler (Qiagen) using the KAPA Library Quantification Kit (Kapa Biosystems, Wilmington, MA). Library integrity and fragment size were determined with a Tapestation 2200 (Agilent Technologies). Sequencing reads were mapped onto the CEN.PK113-7D (Salazar *et al*. 2017) reference genome using the Burrows–Wheeler Alignment (BWA) tool (version 0.7.15) (Li and Durbin 2009) and further processed using SAMtools (version 1.3.1) (Li *et al*. 2009) and Pilon (with -vcf setting; version 1.18) (Walker *et al*. 2014) to identify single nucleotide polymorphisms (SNPs). The sequence was analysed by visualising the generated .bam files in the Integrative Genomics Viewer (IGV) software (version 2.4.0) (Thorvaldsdottir *et al*. 2013). Chromosomal copy number was estimated by the Magnolya algorithm (version 0.15) (Nijkamp *et al*. 2012b).

### Long read sequencing

High quality DNA of IMX2538 was isolated and checked on quantity and quality as described above. Furthermore, quality and integrity of DNA was checked with a TapeStation 2200 (Agilent Technologies, Santa Clara, CA). IMX2538 was sequenced in-house on a single R10 flow cell (FLO—MIN111) using the SQK-LSK109 sequencing kit (Oxford Nanopore Technologies, Oxford, United Kingdom), according to the manufacturer’s instructions. With MinKnow (version 3.6.5, Oxford Nanopore Technologies) raw signal files were generated. Basecalling was performed by Guppy (version 4.0.11, Oxford Nanopore Technologies), followed by *de novo* assembly with Canu (version 2.0 (Koren *et al*. 2017)).

Short- and long-read sequencing data are available at NCBI under BioProject PRJNA757356.

### Growth rate measurement in shake flasks

The growth rate of the constructed strains was determined in 500 mL shake flasks containing 100 mL of SMD or SME medium. Wake-up cultures were inoculated with a 1 mL aliquot of a strain stored at -80 °C and grown until late-exponential phase. Pre-cultures were inoculated from the wake-up cultures and grown to mid-exponential phase. Finally, measuring cultures were inoculated in biological duplicate from the pre-culture at an initial OD_660_ of 0.3. Cultures were monitored for OD_660_ with a Jenway 7200 spectrophotometer in technical duplicate (Cole-Parmer, Vernon Hills, IL). A maximum specific growth rate (µ_max_) was calculated from at least five data points in the exponential phase with at least 2 doublings. Anaerobic shake flask-based experiments were performed at 30 °C in a Bactron anaerobic chamber (Sheldon Manufacturing Inc., Cornelius, OR) with an atmosphere of 5 % (v/v) H_2_, 6 % (v/v) CO_2_ and 89 % (v/v) N_2_, on a IKA KS 260 basic shaker at 200 rpm, using 50 mL shake flasks containing 30 mL SMD (2 %) anaerobic medium.

### Growth rate measurement in microtiter plates

Growth measurements of strains in microtiter plate with a Growth Profiler 960 (EnzyScreen BV, Heemstede, The Netherlands) were performed as described by Postma *et al*. (2021).

### Physiological characterisation of CEN.PK113-7D and IMX2538 in bioreactor cultures

Aerobic batch bioreactor cultures were performed in 2-L bioreactors (Applikon, Delft, The Netherlands). Bioreactors were filled with synthetic medium containing 5.0 g L^-1^ (NH_4_)_2_SO_4_, 3.0 g L^-1^ KH_2_PO_4_, 0.5 g L^-1^ MgSO_4_·7H_2_O, and 1.0 mL L^-1^ trace elements. After heat sterilisation, 20 g L^-1^ glucose, 0.2 g L^-1^ antifoam emulsion C (Sigma-Aldrich, St. Louis, MA), and filter-sterilised vitamins were added to complete the medium. Upon inoculation, bioreactors contained a working volume 1.4 L and the culture pH was maintained at 5.0 by automated addition of 2M H_2_SO_4_ or 2M KOH. Temperature was kept stable at 30°C and mixing of the medium was performed at 800 rpm. The gas flow was set to 700 mL of air per minute to supply oxygen and remove produced carbon dioxide, and an overpressure of 0.3 bar was applied to the reactor. Dissolved oxygen tension was thus maintained for all reactors above 59 % over the whole duration of the batch cultivation. Off-gas was cooled to 2°C in a condenser on the bioreactor to prevent water evaporation, and further dried with a Permapure MD-110-48P-4 filter dryer (Permapure, Lakewood, NJ) for subsequent analysis of carbon dioxide and oxygen percentages by a MultiExact 4100 gas analyser (Servomex, Zoetermeer, The Netherlands). For both CEN.PK113-7D and IMX2538 reactors were run in biological triplicate and inoculated from exponentially growing shake flask cultures.

Optical densities were measured in technical triplicates on Jenway 7200 spectrophotometer (Cole-Parmer) at 660 nm, while cell dry weights were determined by filtration of 10 mL of well-mixed sample over dried PES membrane filters with a pore size of 0.45 µm (Pall Corporation, Port Washington, NY). Filters were washed with demineralized water and dried in a microwave oven for 20 minutes at 360 W. Extracellular organic acids, sugars and ethanol were determined by high performance liquid chromatography (HPLC) analysis using an Aminex HPX-87H ion-exchange column (Agilent, Santa Clara) with 5 mM H_2_SO_4_ as mobile phase and a flow rate of 0.6 mL min^−1^ at 60°C. Glucose, glycerol, and ethanol were detected by a refractive-index detector (Agilent G1362A) and organic acids by a dual-wavelength absorbance detector (Agilent G1314F).

During mid-exponential growth in the glucose consumption phase, intracellular metabolite samples were taken with a filtration-based washing method according to Douma *et al*. (2010) with some modifications. Briefly, approximately 3 mL of cell culture was sampled in 15 mL of 100 % methanol at -40°C. Biomass was washed with cooled 100 % methanol on a PES membrane with a pore size of 0.45 µm (Pall Corporation) which was pre-cooled and wetted with 100 % methanol at -40°C. Finally, metabolites were extracted with 75 % of boiling ethanol. 100 μL of ^13^C cell extract was added to each tube as an internal standard for metabolite quantification (Wu *et al*. 2005). The intracellular CCM metabolites, cofactors and nucleotides were derivatized and quantified as described by de Jonge *et al*. (2011) and Niedenführ *et al*. (2016).

## Abbreviations

CCM: Central Carbon Metabolism
WGD: Whole Genome Duplication
SSD: Smaller-Scale Duplication
MG: Minimal Glycolysis
PPP: Pentose Phosphate Pathway
MC: Mitochondrial Carrier
TCA: cycle Tricarboxylic Acid cycle
glyc_min_: minimized glycolysis
fer_min_: minimized ethanolic fermentation
ppp_min_: minimized pentose phosphate pathway
tca_min_: minimized tricarboxylic acid cycle
mc_min_: minimized mitochondrial carriers
fum_min_: minimized fumarate reductases
glyox_min_: minimized glyoxylate cycle
ace_min_: minimized Acetyl-CoA synthesis
glycerol_min_: minimized glycerol synthesis
Glc: Glucose
Glc-6P: Glucose-6-phosphate
Fru-6P: Fructose-6-phosphate
Fru-1,6-BP: Fructose-1,6-bisphosphate
GAP: Glyceraldehyde-3-phosphate
DHAP: Dihydroxyacetone phosphate
1,3-BPG: 1,3-bisphosphoglycerate
3-PG: 3-phosphoglycerate
2-PG: 2-phosphoglycerate
PEP: Phosphoenolpyruvate
Pyr: Pyruvate
AcAl: Acetaldehyde
EtOH: Ethanol
6P-GLCN-lac: 6-Phosphogluconolactone
6P-GLCN: 6-Phosphoglucononate
RL5P: Ribulose 5-phosphate
R5P: Ribose 5-phosphate
XUL-5P: Xylulose 5-phosphate
S7P: Sedoheptulose 7-phosphate
Ery-4P: Erythrose 4-phosphate
OAA: Oxaloacetate
Cit: Citrate
Cis-Aco: Cis-aconitate
Isocit: Isocitrate
α-KG: α-ketoglutarate
Suc-CoA: Succinyl-CoA
Suc: Succinate
Fum: Fumarate
Mal: Malate
Suc: Succinate
Glyox: Glyoxylate
Ace: Acetate
Ac-CoA: Acetyl-CoA
Glyc-3P: Glycerol-3-phosphate
YPD: Yeast Peptone Dextrose
SMD: Synthetic Medium Dextrose
SME: Synthetic Medium Ethanol
SMGal: Synthetic Medium Galactose
SMMal: Synthetic Medium Maltose
SMSuc: Synthetic Medium Sucrose
SMFruc: Synthetic Medium Fructose
SMPyr: Synthetic Medium Pyruvate

## Acknowledgements

We thank Pilar de la Torre and Marcel van den Broek for sequencing and bioinformatic analysis, Roel Sarelse, Ilse Pardijs and Lycka Kamoen for strain construction, Koen Verhagen, Patricia van Dam and Martin Pabst for analysis of intracellular metabolites, Erik de Hulster for assistance with sampling of the bioreactors, Marijke Luttik for the growth experiment on pyruvate medium, Sofia Dashko and Jean-Marc Daran for input during the literature analysis and Jack Pronk for feedback on an advanced version of the manuscript. This work was funded by the AdLibYeast ERC consolidator grant number 648141 attributed to P.D.-L.

