## Supplementary Material for "Top-down, knowledge-based genetic reduction of yeast central carbon metabolism"

#### Figure S1 - Growth rate of the *OAC1* deletion mutant

Maximum specific growth rate of S. *cerevisiae* CEN.PK113-7D (naïve reference strain) and the *OAC1* deletion mutant, *S. cerevisiae* IMK588, tested in shake flasks on selective SMD medium. Growth rates represent the average and standard deviation of two biological duplicates. * IMK588 had a 26% slower growth rate with respect to CEN.PK113-7D (two-tailed paired homoscedastic t-test p<0.05).

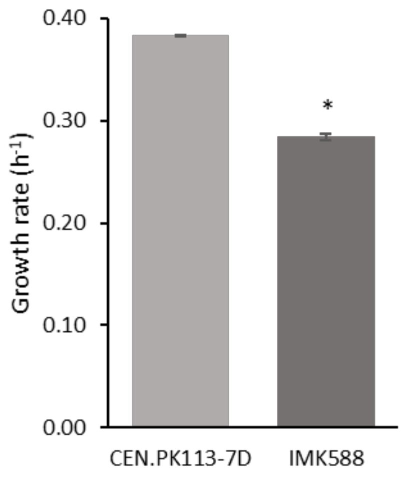

#### Figure S2 - Growth rates of deletion strains

Maximum specific growth rates of all *S. cerevisiae* strains tested in shake flasks on selective SMD (A, B) or SME (B, D) supplemented with uracil. Represented are deletion strains in naïve background (A, B) and in the engineered background (C,D). Colour coding is according to Figure 2 in the main manuscript. Growth rates represent the average and standard deviation of two biological duplicates. Growth rates of the deletion strains are expressed as % of the control strain IMX581. Significant changes in growth rate with respect to the control strain are indicated with a * (two-tailed paired homoscedastic t-test $p<0.05$).

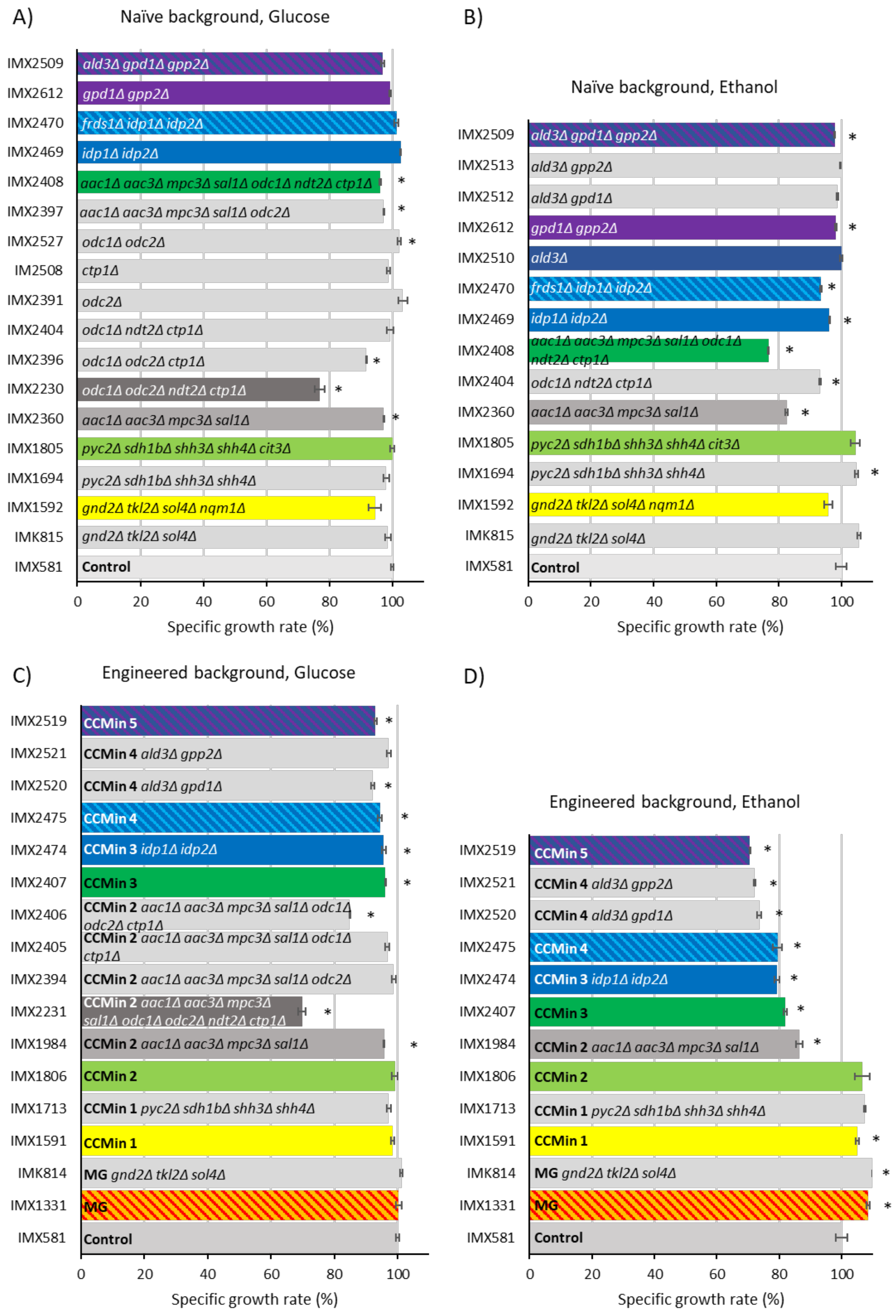
A)
Naïve background, Glucose
IMX2509 ald3Δ gpd1Δ gpp2Δ
IMX2612 gpd1Δ gpp2Δ
IMX2470 frds1Δ idp1Δ idp2Δ
IMX2469 idp1Δ idp2Δ
IMX2408 aac1Δ aac3Δ mpc3Δ sal1Δ odc1Δ ndt2Δ ctp1Δ *
IMX2397 aac1Δ aac3Δ mpc3Δ sal1Δ odc2Δ *
IMX2527 odc1Δ odc2Δ *
IM2508 ctp1Δ
IMX2391 odc2Δ
IMX2404 odc1Δ ndt2Δ ctp1Δ
IMX2396 odc1Δ odc2Δ ctp1Δ *
IMX2230 odc1Δ odc2Δ ndt2Δ ctp1Δ *
IMX2360 aac1Δ aac3Δ mpc3Δ sal1Δ *
IMX1805 pyc2Δ sdh1bΔ shh3Δ shh4Δ cit3Δ
IMX1694 pyc2Δ sdh1bΔ shh3Δ shh4Δ
IMX1592 gnd2Δ tkl2Δ sol4Δ nqm1Δ
IMK815 gnd2Δ tkl2Δ sol4Δ
IMX581 Control
0 20 40 60 80 100
Specific growth rate (%)
B)
Naïve background, Ethanol
IMX2509 ald3Δ gpd1Δ gpp2Δ *
IMX2513 ald3Δ gpp2Δ
IMX2512 ald3Δ gpd1Δ
IMX2612 gpd1Δ gpp2Δ *
IMX2510 ald3Δ
IMX2470 frds1Δ idp1Δ idp2Δ *
IMX2469 idp1Δ idp2Δ *
IMX2408 aac1Δ aac3Δ mpc3Δ sal1Δ odc1Δ ndt2Δ ctp1Δ *
IMX2404 odc1Δ ndt2Δ ctp1Δ *
IMX2360 aac1Δ aac3Δ mpc3Δ sal1Δ *
IMX1805 pyc2Δ sdh1bΔ shh3Δ shh4Δ cit3Δ
IMX1694 pyc2Δ sdh1bΔ shh3Δ shh4Δ *
IMX1592 gnd2Δ tkl2Δ sol4Δ nqm1Δ
IMK815 gnd2Δ tkl2Δ sol4Δ
IMX581 Control
0 20 40 60 80 100
Specific growth rate (%)
C)
Engineered background, Glucose
IMX2519 CCMin 5 *
IMX2521 CCMin 4 ald3Δ gpp2Δ
IMX2520 CCMin 4 ald3Δ gpd1Δ *
IMX2475 CCMin 4 *
IMX2474 CCMin 3 idp1Δ idp2Δ *
IMX2407 CCMin 3 *
IMX2406 CCMin 2 aac1Δ aac3Δ mpc3Δ sal1Δ odc1Δ odc2Δ ctp1Δ *
IMX2405 CCMin 2 aac1Δ aac3Δ mpc3Δ sal1Δ odc1Δ ctp1Δ
IMX2394 CCMin 2 aac1Δ aac3Δ mpc3Δ sal1Δ odc2Δ
IMX2231 CCMin 2 aac1Δ aac3Δ mpc3Δ sal1Δ odc1Δ odc2Δ ndt2Δ ctp1Δ *
IMX1984 CCMin 2 aac1Δ aac3Δ mpc3Δ sal1Δ *
IMX1806 CCMin 2
IMX1713 CCMin 1 pyc2Δ sdh1bΔ shh3Δ shh4Δ
IMX1591 CCMin 1
IMK814 MG gnd2Δ tkl2Δ sol4Δ
IMX1331 MG
IMX581 Control
0 20 40 60 80 100
Specific growth rate (%)
D)
Engineered background, Ethanol
IMX2519 CCMin 5 *
IMX2521 CCMin 4 ald3Δ gpp2Δ *
IMX2520 CCMin 4 ald3Δ gpd1Δ *
IMX2475 CCMin 4 *
IMX2474 CCMin 3 idp1Δ idp2Δ *
IMX2407 CCMin 3 *
IMX1984 CCMin 2 aac1Δ aac3Δ mpc3Δ sal1Δ *
IMX1806 CCMin 2
IMX1713 CCMin 1 pyc2Δ sdh1bΔ shh3Δ shh4Δ
IMX1591 CCMin 1 *
IMK814 MG gnd2Δ tkl2Δ sol4Δ *
IMX1331 MG *
IMX581 Control
0 20 40 60 80 100
Specific growth rate (%)

### Figure S3 - Growth rates in specific environments

A, B, C and D) Specific growth rate determination on osmotic stress conditions (supplemented with uracil) measured with the growth profiler in biological triplicate. E) Specific growth rate of *FRDS1* and *AAC3* deletion mutants on SMD supplemented with Tween, ergosterol and uracil measured under anaerobic conditions in shake flasks in biological duplicate. F) Specific growth rate of *MPC3* complementation strains on SME supplemented with uracil in shake flask in biological duplicate. G) Specific growth rate of *MPC3* complementation strain on SM with 83.3 mM pyruvate supplemented with uracil in shake flask in biological duplicate. Significant changes in growth rate with respect to the parental strain are indicated with a * (two-tailed paired homoscedastic t-test p<0.05).

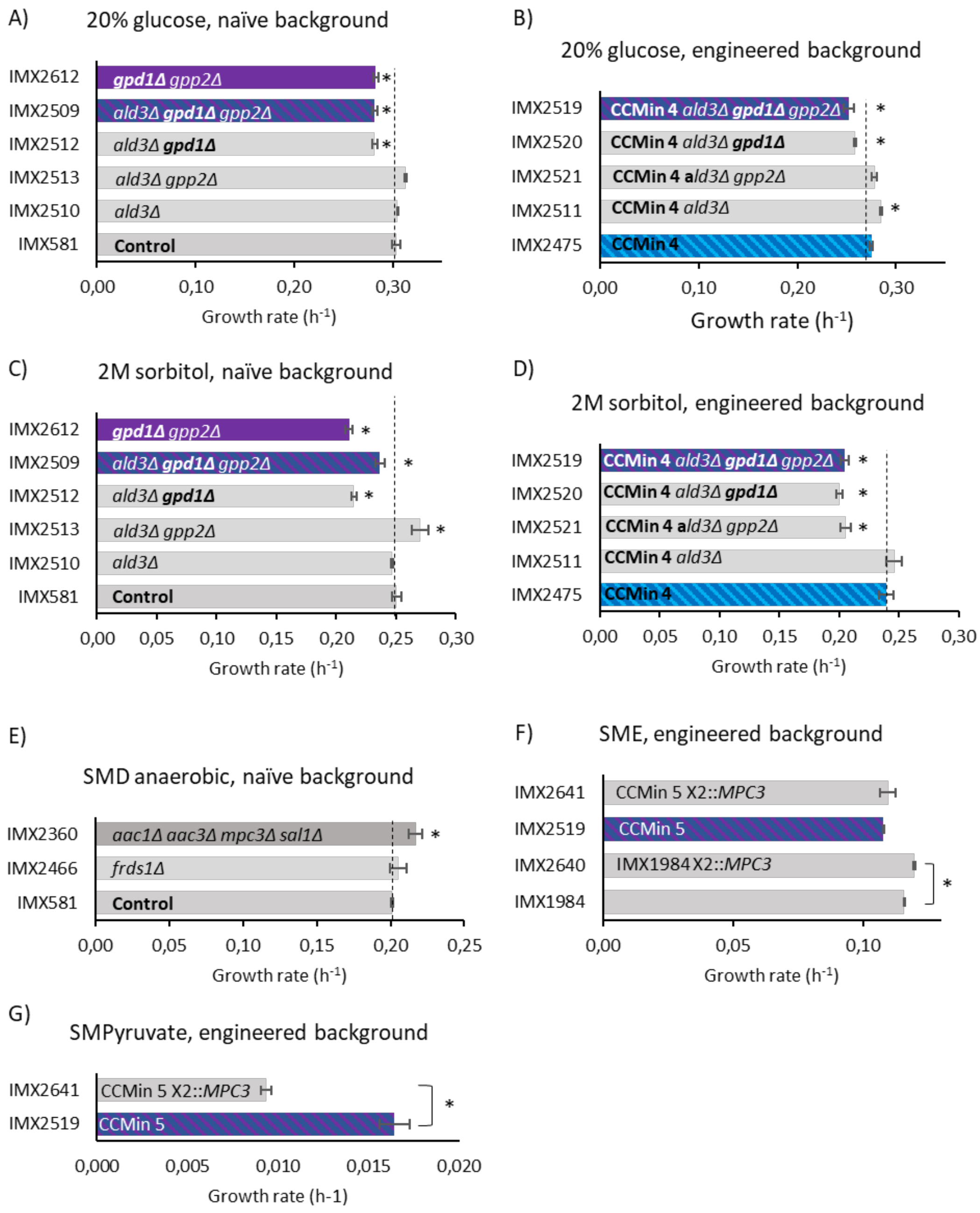

#### Table S1 Genetic characteristics of the paralogs considered for deletion

Genes that were deleted in this study are marked in red. Protein similarity was based on global alignment using BLOSUM62 of the translated genes of the *S. cerevisiae* CEN.PK113-7D genome sequence (Salazar *et al.* 2017). WGD pairs are indicated (Byrne and Wolfe 2005).

| Pathway | Gene | % Protein Similarity | Type of duplication |
|---|---|---|---|
| **Pentose phosphate pathway** | *GND1* (YHR183W) / *GND2* (YGR256W) | 87% | WGD |
| | *TKL1* (YPR074C) / *TKL2* (YBR117C) | 71% | WGD |
| | *SOL3* (YHR163W) / *SOL4* (YGR248W) | 47% | WGD |
| | *TAL1* (YLR354C) / *NQM1* (YGR043C) | 68% | WGD |
| **TCA cycle + anaplerotic reactions** | *PYC1* (YGL062W) / *PYC2* (YBR218C) | 92% | WGD |
| | *SDH1* (YKL148C) / *SDH1b* (YJL045W) | 84% | WGD |
| | *SDH3* (YKL141W) / *SHH3* (YMR118C) | 42% | WGD |
| | *SDH4* (YDR178W) / *SHH4* (YLR164W) | 47% | WGD |
| | *CIT1* (YNR001C) / *CIT2* (YCR005C) / *CIT3* (YPR001W) | CIT1/CIT2=75% **CIT1/CIT3=42%** | CIT1/CIT2=WGD |
| **Mitochondrial transporters** | *AAC1* (YMR056C)/ *AAC2* (YBL030C) / *AAC3* (YBR085W) | AAC2/AAC1=73% AAC2/AAC3=87% | AAC2/AAC3=WGD |
| | *SAL1* (YNL083W) | | |
| | *MPC1* (YGL080W) / *MPC2* (YHR162W) / *MPC3* (YGR243W) | **MPC2/MPC3=65%** MPC2/MPC1=25% | MPC2/MPC3=WGD |
| | *CTP1* (YBR291C) | | |
| | *ODC1* (YPL134C) / *ODC2* (YOR222W) | 61% | WGD |
| | *NDT1* (YIL006W) / *NDT2* (YEL006W) | 51% | WGD |
| **Fumarate reductase** | *OSM1* (YJR051W) / *FRDS1* (YEL047C) | 59% | WGD |
| **Glyoxylate and TCA cycle** | *IDP1* (YDL066W) / *IDP2* (YLR174W) / *IDP3* (YNL009W) | IDP1/IDP3=65% IDP2/IDP3 =73% | IDP2/IDP3=WGD |

| | | | |
|---|---|---|---|
| **acetyl-CoA synthesis** | *ALD2* (YMR170C) / *ALD3* (YMR169C) | 91% | |
| **Glycerol synthesis** | *GPD1* (YDL022W) / *GPD2* (YOL059W) | 72% | WGD |
| | *GPP1* (YIL053W) / *GPP2* (YER062C) | 92% | WGD |

### Table S2 Mitochondrial carrier proteins

A list of 35 mitochondrial transporter identified by Palmieri *et al.* (1996) and reviewed by Palmieri and Monné (2016). In this list the systematic name, standard name, importance in this study and the function of the gene encoding for mitochondrial transporters are listed. NA, not assigned. **CRC1* was not considered as involved in the CCM since there is no carnitine supplied to the cultures in this study and thus the carnitine shuttle is not active.

| Systematic name | Standard name | Considered as involved in CCM? | Function |
|---|---|---|---|
| **YIL006w** | *NDT1* | YES | Mitochondrial $NAD^+$ transporter |
| **YEL006w** | *NDT2* | YES | Mitochondrial $NAD^+$ transporter |
| **YIL134w** | *FLX1* | YES | Mitochondrial FAD transporter |
| **YBR192w** | *RIM2* | NO | Mitochondrial pyrimidine nucleotide transporter |
| **YDL119c** | *HEM25* | NO | Mitochondrial glycine transporter |
| **YNL003c** | *SAM5* | NO | S-adenosylmethionine transporter of the mitochondrial inner membrane |
| **YKR052c** | *MRS4* | NO | Iron transporter of the mitochondrial carrier family (may transport other cations) |
| **YJL133w** | *MRS3* | NO | Iron transporter of the mitochondrial carrier family (May transport other cations) |
| **YGR257c** | *MTM1* | NO | pyridoxal 5'-phosphate (PLP) transporter |
| **YER053c** | *PIC2* | NO | Mitochondrial copper and phosphate carrier |
| **YJR077c** | *MIR1* | NO | Mitochondrial phosphate carrier |
| **YOR130c** | *ORT1* | NO | Ornithine transporter of the mitochondrial inner membrane (ornithine-proton exchange or ornithine-ornithine exchange also transports arginine and lysine) |
| **YOR100c** | *CRC1* | NO* | Mitochondrial inner membrane carnitine transporter. Transports carnitine, acetylcarnitin and propionylcarnitine . |
| **YBR104w** | *YMC2* | NO | Putative mitochondrial inner membrane transporter. Proposed role in oleate metabolism and glutamate biosynthesis. |
| **YPR058w** | *YMC1* | NO | Secondary mitochondrial inner membrane glycine transporter; required with HEM25 for the transport of glycine into mitochondria. Proposed role in oleate metabolism and glutamate biosynthesis. |
| **YPL134c** | *ODC1* | YES | 2-oxodicarboxylate transporter (transports 2-oxoglutarate and oxodipate+ corresponding dicarboxylates and malate by counterexchange) |
| **YOR222w** | *ODC2* | YES | 2-oxodicarboxylate transporter (transports oxogluterate and oxodipate+ corresponding dicarboxylates and malate by counterexchange) |
| **YPR021c** | *AGC1* | NO | Mitochondrial amino acid transporter (transport aspartate and glutamate in uniport as well as in exchange mechanism) |
| **YJR095w** | *SFC1* | YES | Mitochondrial succinate-fumarate counter exchange transporter (fumerate to cytosol, succinate to mitochondria) |
| **YBR291c** | *CTP1* | YES | Mitochondrial inner membrane citrate transporter |
| **YFR045w** | NA | NO | Putative mitochondrial transport protein; null mutant is viable |
| **YMR241w** | *YHM2* | YES | Citrate and oxoglutarate carrier protein (Citrate exported and oxogluterate imported) (oxaloacetate, succinate and fumerate to a lesser extend) |

| | | | |
|---|---|---|---|
| **YLR348c** | *DIC1* | YES | Mitochondrial dicarboxylate carrier (transports malate, succinate, malonate, inorganic phosphate by counter exchange mechanism. Also sulphate, thiosulphate) |
| **YKL120w** | *OAC1* | YES | Transports oxaloacetate and sulfate (unidirectional+counterexchange) |
| **YDL198c** | *GGC1* | NO | Mitochondrial GTP/GDP exchange transporter, essential for mitochondrial genome maintenance, has a role in mitochondrial iron transport |
| **YGR096w** | *TPC1* | YES | Mediates uptake of the essential cofactor thiamine pyrophosphate (ThPP) into mitochondria |
| **YMR056c** | *AAC1* | YES | Mitochondrial inner membrane ADP/ATP translocator |
| **YBL030c** | *AAC2* | YES | Mitochondrial inner membrane ADP/ATP translocator |
| **YBR085w** | *AAC3* | YES | Mitochondrial inner membrane ADP/ATP translocator |
| **YHR002w** | *LEU5* | YES | Involved in the accumulation of CoA in the mitochondrial matrix |
| **YPR011c** | NA | NO | Putative 5'-phosphosulfate (APS) and 3'-phospho-adenosine 5'-phosphosulfate (PAPS) transporter. Not enough information on the function to include in this study. |
| **YNL083w** | *SAL1* | YES | ADP/ATP transporter (activity of either Sal1p or Pet9p is critical for viability) |
| **YGL080w** | *MPC1* | YES | Conserved subunit of mitochondrial pyruvate carrier (MPC) |
| **YHR162w** | *MPC2* | YES | Highly conserved subunit of the mitochondrial pyruvate carrier (MPC) |
| **YGR243w** | *MPC3* | YES | Highly conserved subunit of the mitochondrial pyruvate carrier (MPC) |
| **YMR166C** | *MME1* | NO | Mitochondrial inner membrane transporter that exports magnesium |

#### Table S3 Strains

| Name (Accession no.) | Relevant genotype | Reduced pathway | Origin |
|---|---|---|---|
| | **Unreduced background** | | |
| **CEN.PK113-7D** | *MATa ura3-52 can1Δ::cas9-natNT2 TRP1 LEU2 HIS3* | | (Entian and Kötter 2007) |
| **IMX581 (Y40593)** | *MATa ura3-52 can1Δ::cas9-natNT2 TRP1 LEU2 HIS3* | | (Mans *et al.* 2015) |
| **IMK588** | *MATa URA3 HIS3 LEU2 TRP1 OAC1Δ::kanMX* | | This study |
| **IMK815** | *MATa ura3-52 HIS3 LEU2 TRP1 can1Δ::cas9-natNT2 gnd2Δ tkl2Δ sol4Δ* | | This study |
| **IMX1592** | *MATa ura3-52 HIS3 LEU2 TRP1 can1Δ::cas9-natNT2 gnd2Δ tkl2Δ sol4Δ nqm1Δ* | Pentose phosphate pathway | This study |
| **IMX1694** | *MATa ura3-52 can1Δ::cas9-natNT2 TRP1 LEU2 HIS3 pyc2Δ sdh1bΔ shh3Δ shh4Δ* | | This study |
| **IMX1805** | *MATa ura3-52 can1Δ::cas9-natNT2 TRP1 LEU2 HIS3 pyc2Δ sdh1bΔ shh3Δ shh4Δ cit3Δ* | Anaplerotic reaction and Krebs cycle | This study |
| **IMX2230** | *MATa ura3-52 HIS3 LEU2 TRP1 MAL2-8c SUC2 can1::CAS9-nat odc1Δ odc2Δ ndt2Δ ctp1Δ* | | This study |
| **IMX2360** | *MATa ura3-52 HIS3 LEU2 TRP1 MAL2-8c SUC2 can1::CAS9-nat aac1Δ aac3Δ sal1Δ mpc3Δ* | | This study |
| **IMX2391** | *MATa ura3-52 HIS3 LEU2 TRP1 MAL2-8c SUC2 can1::CAS9-nat odc2Δ* | | This study |
| **IMX2396** | *MATa ura3-52 HIS3 LEU2 TRP1 MAL2-8c SUC2 can1::CAS9-nat ctp1Δ odc1Δ odc2Δ* | | This study |
| **IMX2397** | *MATa ura3-52 HIS3 LEU2 TRP1 MAL2-8c SUC2 can1::CAS9-nat aac1Δ aac3Δ mpc3Δ sal1Δ odc2Δ* | | This study |
| **IMX2404** | *MATa ura3-52 HIS3 LEU2 TRP1 MAL2-8c SUC2 can1::CAS9-nat ctp1Δ odc1Δ ndt2Δ* | | This study |
| **IMX2408** | *MATa ura3-52 HIS3 LEU2 TRP1 MAL2-8c SUC2 can1::CAS9-nat aac1Δ aac3Δ mpc3Δ sal1Δ ctp1Δ odc1Δ ndt2Δ* | Mitochondrial transporters | This study |
| **IMX2416** | *MATa ura3-52 HIS3 LEU2 TRP1 MAL2-8c SUC2 can1::CAS9-nat aac1Δ aac3Δ mpc3Δ sal1Δ ctp1Δ odc1Δ odc2Δ* | | This study |
| **IMX2508** | *MATa ura3-52 HIS3 LEU2 TRP1 MAL2-8c SUC2 can1::CAS9-nat ctp1Δ* | | This study |
| **IMX2527** | *MATa ura3-52 HIS3 LEU2 TRP1 MAL2-8c SUC2 can1::CAS9-nat odc1Δ odc2Δ* | | This study |
| **IMX2466** | *MATa ura3-52 HIS3 LEU2 TRP1 MAL2-8c SUC2 can1::CAS9-nat frds1Δ* | Fumarate reductase | This study |
| **IMX2467** | *MATa ura3-52 HIS3 LEU2 TRP1 MAL2-8c SUC2 can1::CAS9-nat idp1Δ* | | This study |

| | | | |
|---|---|---|---|
| **IMX2468** | *MATa ura3-52 HIS3 LEU2 TRP1 MAL2-8c SUC2 can1::CAS9-nat idp2Δ* | | This study |
| **IMX2469** | *MATa ura3-52 HIS3 LEU2 TRP1 MAL2-8c SUC2 can1::CAS9-nat idp1Δ idp2Δ* | Glyoxylate cycle | This study |
| **IMX2470** | *MATa ura3-52 HIS3 LEU2 TRP1 MAL2-8c SUC2 can1::CAS9-nat frds1Δ idp1Δ idp2Δ* | | This study |
| **IMX2509** | *MATa ura3-52 HIS3 LEU2 TRP1 MAL2-8c SUC2 can1::CAS9-nat ald3Δ gpd1Δ gpp2Δ* | | This study |
| **IMX2510** | *MATa ura3-52 HIS3 LEU2 TRP1 MAL2-8c SUC2 can1::CAS9-nat ald3Δ* | Acetyl-CoA synthesis | This study |
| **IMX2512** | *MATa ura3-52 HIS3 LEU2 TRP1 MAL2-8c SUC2 can1::CAS9-nat ald3Δ gpd1Δ* | | This study |
| **IMX2513** | *MATa ura3-52 HIS3 LEU2 TRP1 MAL2-8c SUC2 can1::CAS9-nat ald3Δ gpp2Δ* | | This study |
| **IMX2612** | *MATa ura3-52 HIS3 LEU2 TRP1 MAL2-8c SUC2 can1::CAS9-nat gpd1Δ gpp2Δ* | Glycerol synthesis | This study |
| | **Reduced background** | | |
| **IMX370** | *MATa ura3-52 his3-1 leu2-3,112 MAL2-8c SUC2 glk1::HIS5, hxk1::LEU2, tdh1, tdh2::AB, gpm2::LoxP, gpm3, eno1, pyk2, pdc5, pdc6, adh2, adh5, adh4* | Glycolysis | (Solis-Escalante *et al.* 2015) |
| **IMX1331 (MG)** | *MATa ura3-52 his3-1 leu2-3,112 MAL2-8c SUC2 glk1::Sphis5 hxk1::KlLEU2 tdh1 tdh2 gpm2 gpm3 eno1 pyk2 pdc5 pdc6 adh2 adh5 adh4 can1Δ::cas9-natNT2* | Glycolysis | This study |
| **IMK814** | *MATa ura3-52 his3-1 leu2-3,112 MAL2-8c SUC2 glk1::Sphis5 hxk1::KlLEU2 tdh1Δ tdh2Δ gpm2Δ gpm3Δ eno1Δ pyk2Δ pdc5Δ pdc6Δ adh2Δ adh5Δ adh4Δ can1Δ::cas9-natNT2 gnd2Δ tkl2Δ sol4Δ* | | This study |
| **IMX1591 (CCMin 1)** | *MATa ura3-52 his3-1 leu2-3,112 MAL2-8c SUC2 glk1::Sphis5 hxk1::KlLEU2 tdh1Δ tdh2Δ gpm2Δ gpm3Δ eno1Δ pyk2Δ pdc5Δ pdc6Δ adh2Δ adh5Δ adh4Δ can1Δ::cas9-natNT2 gnd2Δ tkl2Δ sol4Δ nqm1Δ* | Pentose phosphate pathway | This study |
| **IMX1713** | *MATa ura3-52 his3-1 leu2-3,112 MAL2-8c SUC2 glk1::Sphis5 hxk1::KlLEU2 tdh1Δ tdh2Δ gpm2Δ gpm3Δ eno1Δ pyk2Δ pdc5Δ pdc6Δ adh2Δ adh5Δ adh4Δ can1Δ::cas9-natNT2 gnd2Δ tkl2Δ sol4Δ nqm1Δ pyc2Δ sdh1bΔ shh3Δ shh4Δ* | | This study |
| **IMX1806 (CCMin 2)** | *MATa ura3-52 his3-1 leu2-3,112 MAL2-8c SUC2 glk1::Sphis5 hxk1::KlLEU2 tdh1Δ tdh2Δ gpm2Δ gpm3Δ eno1Δ pyk2Δ pdc5Δ pdc6Δ adh2Δ adh5Δ adh4Δ can1Δ::cas9-natNT2 gnd2Δ tkl2Δ sol4Δ nqm1Δ pyc2Δ sdh1bΔ shh3Δ shh4Δ cit3Δ* | Anaplerotic reaction and Krebs cycle | This study |
| **IMX1984** | *MATa ura3-52 his3-1 leu2-3,112 MAL2-8c SUC2 glk1::Sphis5 hxk1::KlLEU2 tdh1Δ tdh2Δ gpm2Δ gpm3Δ eno1Δ pyk2Δ pdc5Δ pdc6Δ adh2Δ adh5Δ* | | This study |

| | | | |
|---|---|---|---|
| | *adh4Δ can1Δ::cas9-natNT2 gnd2Δ tkl2Δ sol4Δ nqm1Δ pyc2Δ sdh1bΔ shh3Δ shh4Δ cit3Δ aac1Δ aac3Δ sal1Δ mpc3Δ* | | |
| **IMX2231** | *MATa ura3-52 his3-1 leu2-3,112 MAL2-8c SUC2 glk1::Sphis5 hxk1::KlLEU2 tdh1Δ tdh2Δ gpm2Δ gpm3Δ eno1Δ pyk2Δ pdc5Δ pdc6Δ adh2Δ adh5Δ adh4Δ can1Δ::cas9-natNT2 gnd2Δ tkl2Δ sol4Δ nqm1Δ pyc2Δ sdh1bΔ shh3Δ shh4Δ cit3Δ aac1Δ aac3Δ sal1Δ mpc3Δ odc1Δ odc2Δ ndt2Δ ctp1Δ* | | This study |
| **IMX2394** | *MATa ura3-52 his3-1 leu2-3,112 MAL2-8c SUC2 glk1::Sphis5 hxk1::KlLEU2 tdh1Δ tdh2Δ gpm2Δ gpm3Δ eno1Δ pyk2Δ pdc5Δ pdc6Δ adh2Δ adh5Δ adh4Δ can1Δ::cas9-natNT2 gnd2Δ tkl2Δ sol4Δ nqm1Δ pyc2Δ sdh1bΔ shh3Δ shh4Δ cit3Δ aac1Δ aac3Δ sal1Δ mpc3Δ odc2Δ* | | This study |
| **IMX2405** | *MATa ura3-52 his3-1 leu2-3,112 MAL2-8c SUC2 glk1::Sphis5 hxk1::KlLEU2 tdh1Δ tdh2Δ gpm2Δ gpm3Δ eno1Δ pyk2Δ pdc5Δ pdc6Δ adh2Δ adh5Δ adh4Δ can1Δ::cas9-natNT2 gnd2Δ tkl2Δ sol4Δ nqm1Δ pyc2Δ sdh1bΔ shh3Δ shh4Δ cit3Δ aac1Δ aac3Δ sal1Δ mpc3Δ ctp1Δ odc1Δ* | | This study |
| **IMX2406** | *MATa ura3-52 his3-1 leu2-3,112 MAL2-8c SUC2 glk1::Sphis5 hxk1::KlLEU2 tdh1Δ tdh2Δ gpm2Δ gpm3Δ eno1Δ pyk2Δ pdc5Δ pdc6Δ adh2Δ adh5Δ adh4Δ can1Δ::cas9-natNT2 gnd2Δ tkl2Δ sol4Δ nqm1Δ pyc2Δ sdh1bΔ shh3Δ shh4Δ cit3Δ aac1Δ aac3Δ sal1Δ mpc3Δ ctp1Δ odc1Δ odc2Δ* | | This study |
| **IMX2407 (CCMin 3)** | *MATa ura3-52 his3-1 leu2-3,112 MAL2-8c SUC2 glk1Δ::Sphis5 hxk1Δ::KlLEU2 tdh1Δ tdh2Δ gpm2Δ gpm3Δ eno1Δ pyk2Δ pdc5Δ pdc6Δ adh2Δ adh5Δ adh4Δ can1Δ::cas9-natNT2 gnd2Δ tkl2Δ sol4Δ nqm1Δ pyc2Δ sdh1bΔ shh3Δ shh4Δ cit3Δ aac1Δ aac3Δ sal1Δ mpc3Δ ctp1Δ odc1Δ ndt2Δ* | Mitochondrial transporters | This study |
| **IMX2471** | *MATa ura3-52 his3-1 leu2-3,112 MAL2-8c SUC2 glk1Δ::Sphis5 hxk1Δ::KlLEU2 tdh1Δ tdh2Δ gpm2Δ gpm3Δ eno1Δ pyk2Δ pdc5Δ pdc6Δ adh2Δ adh5Δ adh4Δ can1Δ::cas9-natNT2 gnd2Δ tkl2Δ sol4Δ nqm1Δ pyc2Δ sdh1bΔ shh3Δ shh4Δ cit3Δ aac1Δ aac3Δ sal1Δ mpc3Δ ctp1Δ odc1Δ ndt2Δ frds1Δ* | Fumarate reductase | This study |
| **IMX2472** | *MATa ura3-52 his3-1 leu2-3,112 MAL2-8c SUC2 glk1Δ::Sphis5 hxk1Δ::KlLEU2 tdh1Δ tdh2Δ gpm2Δ gpm3Δ eno1Δ pyk2Δ pdc5Δ pdc6Δ adh2Δ adh5Δ adh4Δ can1Δ::cas9-natNT2 gnd2Δ tkl2Δ sol4Δ nqm1Δ pyc2Δ sdh1bΔ shh3Δ shh4Δ cit3Δ aac1Δ aac3Δ sal1Δ mpc3Δ ctp1Δ odc1Δ ndt2Δ idp1Δ* | | This study |

| | | | |
|---|---|---|---|
| **IMX2473** | *MATa ura3-52 his3-1 leu2-3,112 MAL2-8c SUC2 glk1Δ::Sphis5 hxk1Δ::KlLEU2 tdh1Δ tdh2Δ gpm2Δ gpm3Δ eno1Δ pyk2Δ pdc5Δ pdc6Δ adh2Δ adh5Δ adh4Δ can1Δ::cas9-natNT2 gnd2Δ tkl2Δ sol4Δ nqm1Δ pyc2Δ sdh1bΔ shh3Δ shh4Δ cit3Δ aac1Δ aac3Δ sal1Δ mpc3Δ ctp1Δ odc1Δ ndt2Δ idp2Δ* | | This study |
| **IMX2474** | *MATa ura3-52 his3-1 leu2-3,112 MAL2-8c SUC2 glk1Δ::Sphis5 hxk1Δ::KlLEU2 tdh1Δ tdh2Δ gpm2Δ gpm3Δ eno1Δ pyk2Δ pdc5Δ pdc6Δ adh2Δ adh5Δ adh4Δ can1Δ::cas9-natNT2 gnd2Δ tkl2Δ sol4Δ nqm1Δ pyc2Δ sdh1bΔ shh3Δ shh4Δ cit3Δ aac1Δ aac3Δ sal1Δ mpc3Δ ctp1Δ odc1Δ ndt2Δ idp1Δ idp2Δ* | | This study |
| **IMX2475 (CCMin 4)** | *MATa ura3-52 his3-1 leu2-3,112 MAL2-8c SUC2 glk1Δ::Sphis5 hxk1Δ::KlLEU2 tdh1Δ tdh2Δ gpm2Δ gpm3Δ eno1Δ pyk2Δ pdc5Δ pdc6Δ adh2Δ adh5Δ adh4Δ can1Δ::cas9-natNT2 gnd2Δ tkl2Δ sol4Δ nqm1Δ pyc2Δ sdh1bΔ shh3Δ shh4Δ cit3Δ aac1Δ aac3Δ sal1Δ mpc3Δ ctp1Δ odc1Δ ndt2Δ frds1Δ idp1Δ idp2Δ* | Glyoxylate cycle | This study |
| **IMX2511** | *MATa ura3-52 his3-1 leu2-3,112 MAL2-8c SUC2 glk1Δ::Sphis5 hxk1Δ::KlLEU2 tdh1Δ tdh2Δ gpm2Δ gpm3Δ eno1Δ pyk2Δ pdc5Δ pdc6Δ adh2Δ adh5Δ adh4Δ can1Δ::cas9-natNT2 gnd2Δ tkl2Δ sol4Δ nqm1Δ pyc2Δ sdh1bΔ shh3Δ shh4Δ cit3Δ aac1Δ aac3Δ sal1Δ mpc3Δ ctp1Δ odc1Δ ndt2Δ frds1Δ idp1Δ idp2Δ ald3Δ* | Acetyl-Coa synthesis | This study |
| **IMX2519 (CCMin 5)** | *MATa ura3-52 his3-1 leu2-3,112 MAL2-8c SUC2 glk1Δ::Sphis5 hxk1Δ::KlLEU2 tdh1Δ tdh2Δ gpm2Δ gpm3Δ eno1Δ pyk2Δ pdc5Δ pdc6Δ adh2Δ adh5Δ adh4Δ can1Δ::cas9-natNT2 gnd2Δ tkl2Δ sol4Δ nqm1Δ pyc2Δ sdh1bΔ shh3Δ shh4Δ cit3Δ aac1Δ aac3Δ sal1Δ mpc3Δ ctp1Δ odc1Δ ndt2Δ frds1Δ idp1Δ idp2Δ ald3Δ gpd1Δ gpp2Δ* | Glycerol synthesis | This study |
| **IMX2520** | *MATa ura3-52 his3-1 leu2-3,112 MAL2-8c SUC2 glk1Δ::Sphis5 hxk1Δ::KlLEU2 tdh1Δ tdh2Δ gpm2Δ gpm3Δ eno1Δ pyk2Δ pdc5Δ pdc6Δ adh2Δ adh5Δ adh4Δ can1Δ::cas9-natNT2 gnd2Δ tkl2Δ sol4Δ nqm1Δ pyc2Δ sdh1bΔ shh3Δ shh4Δ cit3Δ aac1Δ aac3Δ sal1Δ mpc3Δ ctp1Δ odc1Δ ndt2Δ frds1Δ idp1Δ idp2Δ ald3Δ gpd1Δ* | | This study |
| **IMX2521** | *MATa ura3-52 his3-1 leu2-3,112 MAL2-8c SUC2 glk1Δ::Sphis5 hxk1Δ::KlLEU2 tdh1Δ tdh2Δ gpm2Δ gpm3Δ eno1Δ pyk2Δ pdc5Δ pdc6Δ adh2Δ adh5Δ adh4Δ can1Δ::cas9-natNT2 gnd2Δ tkl2Δ sol4Δ nqm1Δ pyc2Δ sdh1bΔ shh3Δ shh4Δ* | | This study |

| | | |
|---|---|---|
| | *cit3Δ aac1Δ aac3Δ sal1Δ mpc3Δ ctp1Δ odc1Δ ndt2Δ frds1Δ idp1Δ idp2Δ ald3Δ gpp2Δ* | |
| **IMX2538 (minimal CCM strain)** | *MATa ura3-52 his3-1 leu2-3,112 MAL2-8c SUC2 glk1Δ::Sphis5 hxk1Δ::KlLEU2 tdh1Δ tdh2Δ gpm2Δ gpm3Δ eno1Δ pyk2Δ pdc5Δ pdc6Δ adh2Δ adh5Δ adh4Δ can1Δ::cas9-natNT2 gnd2Δ tkl2Δ sol4Δ nqm1Δ pyc2Δ sdh1bΔ shh3Δ shh4Δ cit3Δ aac1Δ aac3Δ sal1Δ mpc3Δ ctp1Δ odc1Δ ndt2Δ frds1Δ idp1Δ idp2Δ ald3Δ gpd1Δ gpp2Δ::URA3* | This study |
| **IMX2640** | *MATa ura3-52 his3-1 leu2-3,112 MAL2-8c SUC2 glk1::Sphis5 hxk1::KlLEU2 tdh1Δ tdh2Δ gpm2Δ gpm3Δ eno1Δ pyk2Δ pdc5Δ pdc6Δ adh2Δ adh5Δ adh4Δ can1Δ::cas9-natNT2 gnd2Δ tkl2Δ sol4Δ nqm1Δ pyc2Δ sdh1bΔ shh3Δ shh4Δ cit3Δ aac1Δ aac3Δ sal1Δ mpc3Δ X2::pMPC3-MPC3-tMPC3* | This study |
| **IMX2641** | *MATa ura3-52 his3-1 leu2-3,112 MAL2-8c SUC2 glk1Δ::Sphis5 hxk1Δ::KlLEU2 tdh1Δ tdh2Δ gpm2Δ gpm3Δ eno1Δ pyk2Δ pdc5Δ pdc6Δ adh2Δ adh5Δ adh4Δ can1Δ::cas9-natNT2 gnd2Δ tkl2Δ sol4Δ nqm1Δ pyc2Δ sdh1bΔ shh3Δ shh4Δ cit3Δ aac1Δ aac3Δ sal1Δ mpc3Δ ctp1Δ odc1Δ ndt2Δ frds1Δ idp1Δ idp2Δ ald3Δ gpd1Δ gpp2Δ X2::pMPC3-MPC3-tMPC3* | This study |

#### Table S4 Plasmids

N.A., not applicable.

| Name | Relevant characteristics | Primer(s) used for gRNA | Origin |
|---|---|---|---|
| **p414-*TEF1p-Cas9-CYC1t*** | CEN6/ARS ampR *TEF1p-Cas9-CYC1t* | N.A. | (DiCarlo *et al.* 2013) |
| **pUG-*natNT2*** | ampR *AgTEF1p-nat-AgTEF1t* | N.A. | (de Kok *et al.* 2012) |
| **pUG6** | ampR *AgTEF1p-kanMX-AgTEF1t* | N.A. | (Güldener *et al.* 1996) |
| **pMEL10** | 2µm ampR *KlURA3* gRNA-*CAN1*.Y | N.A. | (Mans *et al.* 2015) |
| **pMEL11** | 2µm ampR *amdSYM* gRNA-*CAN1*.Y | N.A | (Mans *et al.* 2015) |
| **pROS10** | 2µm ampR *URA3* gRNA-*CAN1*.Y gRNA-*ADE2*.Y | N.A | (Mans *et al.* 2015) |
| **pROS11** | 2µm ampR *amdSYM* gRNA-*CAN1*.Y gRNA-*ADE2*.Y | N.A | (Mans *et al.* 2015) |
| **pUDR286** | 2µm ampR *URA3* gRNA-*TKL2* gRNA-*SOL4* | 9508 & 9503 | This study |
| **pUDR287** | 2µm ampR *amdSYM* gRNA-*GND2* | 7231 | This study |
| **pUDR351** | 2µm ampR *URA3* gRNA-*CIT3* | 12537 | This study |
| **pUDR353** | 2µm ampR *URA3* gRNA-*NQM1* | 12569 | This study |
| **pUDR354** | 2µm ampR *URA3* gRNA-*PYC2* gRNA-*SDH1b* | 12514 & 12521 | This study |
| **pUDR355** | 2µm ampR *amdSYM* gRNA-*SHH3* gRNA-*SHH4* | 9446 & 12529 | This study |
| **pUDR376** | 2µm ampR *amdSYM* gRNA-*X2* (2x) | N.A. | (Wronska *et al.* 2020) |
| **pUDR458** | 2µm ampR *URA3* gRNA-*AAC1* gRNA-*AAC3* | 13820 & 13826 | This study |
| **pUDR460** | 2µm ampR *amdSYM* gRNA-*CTP1* gRNA-*NDT2* | 9489 & 13853 | This study |
| **pUDR462** | 2µm ampR *amdSYM* gRNA-*SAL1* gRNA-*MPC3* | 13830 & 13859 | This study |
| **pUDR606** | 2µm ampR *amdSYM* gRNA-*ODC1* gRNA-*ODC2* | 13840 & 15580 | This study |
| **pUDR686** | 2µm ampR *URA3* gRNA-*ODC1* gRNA-*CTP1* | 13840 & 9489 | This study |
| **pUDR687** | 2µm ampR *amdSYM* gRNA-*NDT2* (2x) | 13853 | This study |
| **pUDR688** | 2µm ampR *amdSYM* gRNA-*ODC2* (2x) | 13846 | This study |
| **pUDR722** | 2µm ampR *amdSYM* gRNA-*FRDS1* (2x) | 17280 | This study |
| **pUDR723** | 2µm ampR *URA3* gRNA-*IDP1* (2x) | 12285 | This study |
| **pUDR724** | 2µm ampR *URA3* gRNA-*IDP2* (2x) | 17287 | This study |
| **pUDR725** | 2µm ampR *URA3* gRNA-*IDP1* gRNA-*IDP2* | 12285 & 17287 | This study |
| **pUDR738** | 2µm ampR *URA3* gRNA-*CTP1* (2x) | 9489 | This study |
| **pUDR739** | 2µm ampR *URA3* gRNA-*ALD3* (2x) | 17446 | This study |
| **pUDR740** | 2µm ampR *amdSYM* gRNA-*GPD1* gRNA-*GPP2* | 9498 | This study |
| **pUDR741** | 2µm ampR *URA3* gRNA-*GPD1* gRNA-*ALD3* | 7772 & 17446 | This study |
| **pUDR742** | 2µm ampR *URA3* gRNA-*GPP2* gRNA-*ALD3* | 9498 & 17446 | This study |

#### Table S5 pROS/pMEL gRNA primers. gRNA sequence is underlined.

| Primer name | Primer number | Sequence (5’ to 3’) |
|---|---|---|
| **2mu inside fw** | 5974 | TACTTTTGAGCAATGTTTGTGGA |
| **2mu inside rv** | 5975 | AACGAGCTACTAAAATATTGCGAA |
| **Primer_gRNA_426_fw** | 5979 | TATTGACGCCGGGCAAGAGC |
| **Primer_stRNA_426_rv** | 5980 | CGACCGAGTTGCTCTTG |
| **P426 CRISPR rv** | 6005 | GATCATTTATCTTTCACTGCGGAGAAG |
| **RV_gnd2_gRNA** | 7231 | GTTGATAACGGACTAGCCTTATTTTAACTTGCTATTTCTAGCT<br>CTAAAACTATGATCTGGCAGCTTCGCGGATCATTTATCTTTCA<br>CTGCGGAGAAGTTTCGAACGCCGAAACATGCGCA |
| **SOL4_targetRNA FW** | 9503 | TGCGCATGTTTCGGCGTTCGAAACTTCTCCGCAGTGAAAGAT<br>AAATGATCACATTTTTCCACATATTAAGGTTTTAGAGCTAGA<br>AATAGCAAGTTAAAATAAG |
| **TKL2_targetRNA FW** | 9508 | TGCGCATGTTTCGGCGTTCGAAACTTCTCCGCAGTGAAAGAT<br>AAATGATCTCAAAAACTTAATGAGGAATGTTTTAGAGCTAGA<br>AATAGCAAGTTAAAATAAG |
| **SHH3 gRNA fw** | 9446 | TGCGCATGTTTCGGCGTTCGAAACTTCTCCGCAGTGAAAGAT<br>AAATGATCATCCGGTTTTTATAACCCCAGTTTTAGAGCTAGA<br>AATAGCAAGTTAAAATAAGGCTAGTCCGTTATCAAC |
| **CTP1_targetRNA FW** | 9489 | TGCGCATGTTTCGGCGTTCGAAACTTCTCCGCAGTGAAAGAT<br>AAATGATCACACCAAAATACCATAATAAGTTTTAGAGCTAGA<br>AATAGCAAGTTAAAATAAGGCTAGTCCGTTATCAAC |
| **GPP2_targetRNA FW** | 9498 | TGCGCATGTTTCGGCGTTCGAAACTTCTCCGCAGTGAAAGAT<br>AAATGATCCAGAACCATATTTGAAAGGCGTTTTAGAGCTAGA<br>AATAGCAAGTTAAAATAAG |
| **gRNA_IDP1_fwd** | 12285 | TGCGCATGTTTCGGCGTTCGAAACTTCTCCGCAGTGAAAGAT<br>AAATGATCGATCTTATCCCAAATGATACGTTTTAGAGCTAGA<br>AATAGCAAGTTAAAATAAGGCTAGTCCGTTATCAAC |
| **PYC2_targetRNA fw** | 12514 | TGCGCATGTTTCGGCGTTCGAAACTTCTCCGCAGTGAAAGAT<br>AAATGATCGTGACTTAAATAAGAAAACTGTTTTAGAGCTAGA<br>AATAGCAAGTTAAAATAAGGCTAGTCCGTTATCAAC |
| **SDH1b_targetRNA fw** | 12521 | TGCGCATGTTTCGGCGTTCGAAACTTCTCCGCAGTGAAAGAT<br>AAATGATCTTTCTGATAAGTCAATGATCGTTTTAGAGCTAGA<br>AATAGCAAGTTAAAATAAGGCTAGTCCGTTATCAAC |
| **SHH4_targetRNA fw** | 12529 | TGCGCATGTTTCGGCGTTCGAAACTTCTCCGCAGTGAAAGAT<br>AAATGATCCATATAAGAGGAATCATTTGGTTTTAGAGCTAGA<br>AATAGCAAGTTAAAATAAGGCTAGTCCGTTATCAAC |
| **CIT3_targetRNA rv** | 12537 | GTTGATAACGGACTAGCCTTATTTTAACTTGCTATTTCTAGCT<br>CTAAAACCATTTGTAATGTTCGAATAAGATCATTTATCTTTCA<br>CTGCGGAGAAGTTTCGAACGCCGAAACATGCGCA |
| **NQM1_targetRNA RV** | 12569 | GTTGATAACGGACTAGCCTTATTTTAACTTGCTATTTCTAGCT<br>CTAAAACTCTAGAACAGTTATATGAATGATCATTTATCTTTCA<br>CTGCGGAGAAGTTTCGAACGCCGAAACATGCGCA |
| **AAC1_targetRNA FW** | 13820 | TGCGCATGTTTCGGCGTTCGAAACTTCTCCGCAGTGAAAGAT<br>AAATGATCGTATAAGAAGACACTGAAAAGTTTTAGAGCTAG<br>AAATAGCAAGTTAAAATAAGGCTAGTCCGTTATCAAC |

| | | |
|---|---|---|
| **AAC3_targetRNA FW** | 13826 | TGCGCATGTTTCGGCGTTCGAAACTTCTCCGCAGTGAAAGAT AAATGATCTTATAAAAAGACCTTGAAATGTTTTAGAGCTAGA AATAGCAAGTTAAAATAAGGCTAGTCCGTTATCAAC |
| **SAL1_targetRNA FW** | 13830 | TGCGCATGTTTCGGCGTTCGAAACTTCTCCGCAGTGAAAGAT AAATGATCTTGGCATTAACGAAATAAATGTTTTAGAGCTAGA AATAGCAAGTTAAAATAAGGCTAGTCCGTTATCAAC |
| **ODC1_targetRNA FW** | 13840 | TGCGCATGTTTCGGCGTTCGAAACTTCTCCGCAGTGAAAGAT AAATGATCAAGAATAGTGTTGTGAAAGGGTTTTAGAGCTAG AAATAGCAAGTTAAAATAAGGCTAGTCCGTTATCAAC |
| **ODC2_targetRNA FW** | 13846 | TGCGCATGTTTCGGCGTTCGAAACTTCTCCGCAGTGAAAGAT AAATGATCGTGCTGTTAAAAAATACAACGTTTTAGAGCTAGA AATAGCAAGTTAAAATAAGGCTAGTCCGTTATCAAC |
| **ODC2_targetRNA_fw** | 15580 | TGCGCATGTTTCGGCGTTCGAAACTTCTCCGCAGTGAAAGAT AAATGATCTTTTTCAGGGATCTGAAGTAGTTTTAGAGCTAGA AATAGCAAGTTAAAATAAGGCTAGTCCGTTATCAAC |
| **NDT2_targetRNA FW** | 13853 | TGCGCATGTTTCGGCGTTCGAAACTTCTCCGCAGTGAAAGAT AAATGATCGAAAATTTAAAAATAAGGTTGTTTTAGAGCTAGA AATAGCAAGTTAAAATAAGGCTAGTCCGTTATCAAC |
| **MPC3_targetRNA FW** | 13859 | TGCGCATGTTTCGGCGTTCGAAACTTCTCCGCAGTGAAAGAT AAATGATCTTATTCACTACATAATAAAAGTTTTAGAGCTAGA AATAGCAAGTTAAAATAAGGCTAGTCCGTTATCAAC |
| **FRDS1_targetRNA FW** | 17280 | TGCGCATGTTTCGGCGTTCGAAACTTCTCCGCAGTGAAAGAT AAATGATCTAAAGGTGTCCAAGAATTAAGTTTTAGAGCTAGA AATAGCAAGTTAAAATAAGGCTAGTCCGTTATCAAC |
| **IDP1_targetRNA FW** | 12285 | TGCGCATGTTTCGGCGTTCGAAACTTCTCCGCAGTGAAAGAT AAATGATCGATCTTATCCCAAATGATACGTTTTAGAGCTAGA AATAGCAAGTTAAAATAAGGCTAGTCCGTTATCAAC |
| **IDP2_targetRNA FW** | 17287 | TGCGCATGTTTCGGCGTTCGAAACTTCTCCGCAGTGAAAGAT AAATGATCATGAGCAAACAAGAATAATCGTTTTAGAGCTAG AAATAGCAAGTTAAAATAAGGCTAGTCCGTTATCAAC |
| **ALD3_targetRNA FW** | 17446 | TGCGCATGTTTCGGCGTTCGAAACTTCTCCGCAGTGAAAGAT AAATGATCAACTTTAGCTTCTTCTTGATGTTTTAGAGCTAGAA ATAGCAAGTTAAAATAAGGCTAGTCCGTTATCAAC |
| **GPD1_targetRNA FW** | 7772 | TGCGCATGTTTCGGCGTTCGAAACTTCTCCGCAGTGAAAGAT AAATGATCATTCTTCAATCATGTCCGGCGTTTTAGAGCTAGA AATAGCAAGTTAAAATAAG |

#### Table S6 List of primers used to confirm guideRNA plasmids

| Primer name | Primer number | Sequence (5' to 3') |
|---|---|---|
| **FK105-MP1** | 2528 | TCTTTCCTGCGTTATCCC |
| **Fus Tag B fw** | 4672 | CACCTTTCGAGAGGACGATG |
| **LP crRNA rv** | 5941 | GCTGGCCTTTTGCTCACATG |
| **RV_gnd2_gRNA_check** | 7257 | TATGATCTGGCAGCTTCGCG |
| **TKL2_beta_dg rv** | 9708 | ATTCCTCATTAAGTTTTTGA |
| **SOL4_alpha_dg rv** | 9709 | CTTAATATGTGGAAAAATGT |
| **GPP2_beta_dg rv** | 9710 | GCCTTTCAAATATGGTTCTG |
| **TEFterm Fw** | 9719 | TCAAGAACTTGTCATTTGTATAG |
| **p426_AmdS_origin_removal_2** | 10466 | GGAAATGTGCGCGGAAC |
| **fw_CYC1t** | 11787 | TCATGTAATTAGTTATGTCACGC |
| **PYC2_pROS_dg rv** | 12520 | AGTTTTCTTATTTAAGTCAC |
| **SDH1b_pROS_dg rv** | 12527 | GATCATTGACTTATCAGAAA |
| **SHH3_pROS_dg rv** | 12528 | TGGGGTTATAAAAACCGGAT |
| **SHH4_pROS_dg rv** | 12535 | CAAATGATTCCTCTTATATG |
| **CIT3_pMEL_dg rv** | 12542 | TTATTCGAACATTACAAATG |
| **NQM1_pMEL_dg fw** | 12729 | ATTCATATAACTGTTCTAGA |
| **AAC1_pROS_dg rv** | 13825 | TTTTCAGTGTCTTCTTATAC |
| **AAC3_pROS_dg rv** | 13829 | ATTTCAAGGTCTTTTTATAA |
| **SAL1_pROS_dg rv** | 13835 | ATTTATTTCGTTAATGCCAA |
| **ODC1_pROS_dg rv** | 13845 | CCTTTCACAACACTATTCTT |
| **ODC2_pROS_dg rv** | 13851 | GTTGTATTTTTTAACAGCAC |
| **CTP1_pROS_dg rv** | 13852 | TTATTATGGTATTTTGGTGT |
| **NDT2_pROS_dg rv** | 13858 | AACCTTATTTTTAAATTTTC |
| **MPC3_pMEL_dg fw** | 13864 | TTATTCACTACATAATAAAA |
| **FRDS1_pROS_dg rv** | 17285 | CTTAATTCTTGGACACCTTTAGATC |
| **IDP1_pROS_dg rv** | 17286 | CGTATCATTTGGGATAAGATCGATC |
| **IDP2_pROS_dg rv** | 17290 | CGATTATTCTTGTTTGCTCATGATC |
| **GPD1_pROS_dg rv** | 17443 | TATGGTCGACGTCCTTGCCC |
| **ALD3_pROS_dg rv** | 17447 | CATCAAGAAGAAGCTAAAGTTGATC |

#### Table S7 List of primers used for making repair fragments for gene deletion or integration

| Primer name | Primer number | Sequence (5' to 3') |
|---|---|---|
| **OAC1_KO_fw** | 6358 | ATAGCAAGTCAGACACAAGCACATCTCATCGAATTATATCG TAAGCAAATGCCAGCTGAAGCTTCGTACG |
| **OAC1_KO_rv** | 6359 | CTGGCCAATGAATGAAACTTCAAACCTCGGAGTTTGTTATG GGAATTAATGCATAGGCCACTAGTGGATCTG |
| **Gnd2_repair_FW_new** | 7299 | AAGAATTCGTAGGTGCAGGTGAGCATATTGCCGGATAAGT GTAGTTACGCAACTACAATTGTTACTAAGGCCCAATCCGGT TGGAGAAGAACTATTGCCCTTGCTGCTACTTACGGTATT |
| **Gnd2_repair_RV_new** | 7300 | AATACCGTAAGTAGCAGCAAGGGCAATAGTTCTTCTCCAA CCGGATTGGGCCTTAGTAACAATTGTAGTTGCGTAACTAC ACTTATCCGGCAATATGCTCACCTGCACCTACGAATTCTT |
| **SHH3 repair oligo fw** | 9448 | CTCAATGCGACTGTGATAGCTGATAAGTGGAGCTCAGAAA TATTCAGAAGCGTAAGAATAATGAAATGTAAGGAATGTAC TTAGTGTTTTTTGCTACATAGCTTTTATGATACTCTTTAT |
| **SHH3 repair oligo rv** | 9449 | ATAAAGAGTATCATAAAAGCTATGTAGCAAAAAACACTAA GTACATTCCTTACATTTCATTATTCTTACGCTTCTGAATATT TCTGAGCTCCACTTATCAGCTATCACAGTCGCATTGAG |
| **CTP1_repair oligo fw** | 9491 | TTGATGTCACAATGAAAGAACTCCAAAGTAGAGCTTGAAT TATAAATTAGCATTTTACCGAATGTATTATTGTGTACAATA TATCATCTAATGTTTTCTACTCGTTATAAGTCTATTTAC |
| **CTP1_repair oligo rv** | 9492 | GTAAATAGACTTATAACGAGTAGAAAACATTAGATGATAT ATTGTACACAATAATACATTCGGTAAAATGCTAATTTATAA TTCAAGCTCTACTTTGGAGTTCTTTCATTGTGACATCAA |
| **GPP2_repair oligo fw** | 9499 | GTTTGCCAAAGGTTTCTTTTCTGCTCAATTTGGTCTAACTCT TTTCATATTAATAGCGCCATTAAATAAATACGTAGATAGAT TTTTTTTTTTAAAACATATAGTGTGCTATTATTTCTG |
| **GPP2_repair oligo rv** | 9500 | CAGAAATAATAGCACACTATATGTTTTAAAAAAAAAAATCT ATCTACGTATTTATTTAATGGCGCTATTAATATGAAAAGAG TTAGACCAAATTGAGCAGAAAAGAAACCTTTGGCAAAC |
| **SOL4_repair oligo fw** | 9504 | CAGCAGTTTTCCAAACAAAGAATGCCATTCATCAAATAATC CACAACCACCTCAAGAAAATTACACTCGTCTTTATACGAAA CTGGCTCCGTTAATCACGACAGACAACCTTAATTACAT |
| **SOL4_repair oligo rv** | 9505 | ATGTAATTAAGGTTGTCTGTCGTGATTAACGGAGCCAGTTT CGTATAAAGACGAGTGTAATTTTCTTGAGGTGGTTGTGGA TTATTTGATGAATGGCATTCTTTGTTTGGAAAACTGCTG |
| **TKL2_repair oligo fw** | 9509 | TTGTTGGGAGGAGTCCTGAATAAGGAGTGTCGAATATAG GGAGCTTCATTCGTTGTCAAGGAAGTAAACAGTTCTTTGCT ATTTCACACTTCCTGGTTGATGGTCACTTGCTGCCTGAAA |
| **TKL2_repair oligo rv** | 9510 | TTTCAGGCAGCAAGTGACCATCAACCAGGAAGTGTGAAAT AGCAAAGAACTGTTTACTTCCTTGACAACGAATGAAGCTC CCTATATTCGACACTCCTTATTCAGGACTCCTCCCAACAA |
| **IDP1_repair_fwd** | 12295 | TTATGAAATCTTCCTTCAAGCAATTGTGAGACAACAGACGC ACAAGGAAGATCGCCCAGCTCGAATTTACGTAGCCCAATC TACCACTTTTTTTTTCATTTTTTAAAGTGTTATACTTAG |

| | | |
|---|---|---|
| **IDP1_repair_rev** | 12296 | CTAAGTATAACACTTTAAAAAATGAAAAAAAAAGTGGTAG ATTGGGCTACGTAAATTCGAGCTGGGCGATCTTCCTTGTG CGTCTGTTGTCTCACAATTGCTTGAAGGAAGATTTCATAA |
| **PYC2-repair oligo fw** | 12516 | TGCAAAATAAAGGACAGTTACTAGGAGAGAAAATAAGGG ACATAGAGAACAAAATAAAATTTTTTACTCGTTAATTATAT TTTATGACATCTGAAAATACTAGCTGTACTATATATGGCG |
| **PYC2-repair oligo rv** | 12517 | CGCCATATATAGTACAGCTAGTATTTTCAGATGTCATAAAA TATAATTAACGAGTAAAAAATTTTACTTTGTTCTCTATGTCT CTTATTTTCTCTCCTAGTAACTGTCCTTTATTTTGCA |
| **SDH1b -repair oligo fw** | 12523 | GAAAAGAAGGAGCCTAAATACGTATATCTATATACATGTA TACACGTGAGCTAATAAATTTTCTTATTTATTTATTTATTTA TTTTGGAGGGCAAACTTATTTATTGATCTGGCAAAAAT |
| **SDH1b -repair oligo rv** | 12524 | ATTTTTGCCAGATCAATAAATAAGTTTGCCCTCCAAAATAA ATAAATAAATAAATAAGAAAATTTATTAGCTCACGTGTATA CATGTATATAGATATACGTATTTAGGCTCCTTCTTTTC |
| **SHH4-repair oligo fw** | 12531 | AGTTCTAATGAATCAGCAAAGATTCTCAAAAAGGTTGCCC CAATTCTTAGGAAAGTAGGATCAATATGGTTTGGTTAGTG GTGACTACCTTTTTTATTCTCGTTATATATGTGTATTAGA |
| **SHH4-repair oligo rv** | 12532 | TCTAATACACATATATAACGAGAATAAAAAAGGTAGTCAC CACTAACCAAACCATATTGATCCTACTTTCCTAAGAATTGG GGCAACCTTTTTGAGAATCTTTGCTGATTCATTAGAACT |
| **CIT3-repair oligo fw** | 12538 | AAAAAGATCGTATTTGATCAAGAATTTATACATAGACGCC GCTAAATAATTGAATACAAACGCAGTTCCAATTTACAAGA ATGCTTCGTTTGCTATTACAATATTGAAATATAAATAAAA |
| **CIT3-repair oligo rv** | 12539 | TTTTATTTATATTTCAATATTGTAATAGCAAACGAAGCATTC TTGTAAATTGGAACTGCGTTTGTATTCAATTATTTAGCGGC GTCTATGTATAAATTCTTGATCAAATACGATCTTTTT |
| **NQM1_repair oligo fw** | 12570 | TTCTTGCTAGCGTAAGTCATAAAAAAATAGGAAATAATCAC ATATATACAAGAAATTAAATTCATTAAGAGTAGAGGTACC TACTTATATATATAAATATATATATACCACTTTCCTTTTC |
| **NQM1_repair oligo rv** | 12571 | GAAAAGGAAAGTGGTATATATATATTTATATATATAAGTA GGTACCTCTACTCTTAATGAATTTAATTTCTTGTATATATGT GATTATTTCCTATTTTTTATGACTTACGCTAGCAAGAA |
| **AAC1_repair oligo fw** | 13821 | TTCCTTTTTACAGCAGTAATGTCTCACACAGAAACATAGAC TCAGCAGTCACACTTCGGTAAAAAAAAGAAAACAACAAAC GAATAAAATCTAAAAATTCTACATATTCTTGCTATTTAT |
| **AAC1_repair oligo rv** | 13822 | ATAAATAGCAAGAATATGTAGAATTTTTAGATTTTATTCGT TTGTTGTTTTCTTTTTTTTACCGAAGTGTGACTGCTGAGTCT ATGTTTCTGTGTGAGACATTACTGCTGTAAAAAGGAA |
| **AAC3_repair oligo fw** | 13827 | GTATAATTAACTCAATTGAAGACGGTTTACCTGAAGTGAT ATACTGTGCCTTGAGAAACATCAGTTGGATGAAGAAAAAA GTCATTTTCTCGACTTCTCTTCACCTTTCGATCGATTTGA |
| **AAC3_repair oligo rv** | 13828 | TCAAATCGATCGAAAGGTGAAGAGAAGTCGAGAAAATGA CTTTTTTCTTCATCCAACTGATGTTTCTCAAGGCACAGTATA TCACTTCAGGTAAACCGTCTTCAATTGAGTTAATTATAC |
| **SAL1_repair oligo fw** | 13831 | GGGAAATATTGAATATTTGAAGGGGCAAATAACGTTTAAT TCAATCGAATGACGGTCTAATGATAATAACATATGCATAT GTATCAACTAGTGAATTCTATTTAATTATAAACCGCTGCT |

| | | |
|---|---|---|
| **SAL1_repair oligo rv** | 13832 | AGCAGCGGTTTATAATTAAATAGAATTCACTAGTTGATACA<br>TATGCATATGTTATTATCATTAGACCGTCATTCGATTGAATT<br>AAACGTTATTTGCCCCTTCAAATATTCAATATTTCCC |
| **ODC1_repair oligo fw** | 13841 | TTTATCTCATTTATTCTCAAGATAGAACTGTAACGAGGGTG<br>AAAGAAAAAGAACTGTAAGTCCTGACGCTCTTTATTTCATT<br>TTGTTGTAGCCCGCCCATACATGTATACGTATATATAT |
| **ODC1_repair oligo rv** | 13842 | ATATATATACGTATACATGTATGGGCGGGCTACAACAAAA<br>TGAAATAAAGAGCGTCAGGACTTACAGTTCTTTTTCTTTCA<br>CCCTCGTTACAGTTCTATCTTGAGAATAAATGAGATAAA |
| **ODC2_repair oligo fw** | 13847 | CACTTTTTTACTGGGTTAGATTCTATATAGTCAAGTAATTC<br>GAGGTAGCATCAAAGTAATTATTTATTGTATATAGAATACC<br>TCTTTCCTCTTCAATCTTAACTACGTTATTTCTACGTC |
| **ODC2_repair oligo rv** | 13848 | GACGTAGAAATAACGTAGTTAAGATTGAAGAGGAAAGAG<br>GTATTCTATATACAATAAATAATTACTTTGATGCTACCTCG<br>AATTACTTGACTATATAGAATCTAACCCAGTAAAAAAGTG |
| **NDT2_repair oligo fw** | 13854 | GTTGTTCTCAACTTATTTTCATTGTTTGATGAAAAGCGGAA<br>TATACATCACGGCTATATAAGTGAAACAGAAAAATGTACA<br>TGACACAAGCTGAGGGTCGTAAAAAGCATCTTTCGCAAT |
| **NDT2_repair oligo rv** | 13855 | ATTGCGAAAGATGCTTTTTACGACCCTCAGCTTGTGTCATG<br>TACATTTTTCTGTTTCACTTATATAGCCGTGATGTATATTCC<br>GCTTTTCATCAAACAATGAAAATAAGTTGAGAACAAC |
| **MPC3_repair oligo fw** | 13860 | GTCTTTAAGACTATACGCATAAGCATTCAAGACACATAGA<br>AACACAAACCTATATTTTTATTACGTAAACGATAATATGTT<br>CCTGAACTCGCATTTTTTTAATGATTTTTTATGACCTCT |
| **MPC3_repair oligo rv** | 13861 | AGAGGTCATAAAAAATCATTAAAAAAATGCGAGTTCAGGA<br>ACATATTATCGTTTACGTAATAAAAATATAGGTTTGTGTTT<br>CTATGTGTCTTGAATGCTTATGCGTATAGTCTTAAAGAC |
| **FRDS1_repair oligo fw** | 17281 | CATAGCACCCCATTTTTTTTTCTTATCTTCACTTCAACAACCC<br>CTCATCCGTAATTGTAAGGTGGTGCTAGAATCAATGTCAA<br>GGCTCAAGTCATTGGCAAGAACGACGAAAGGCTACTA |
| **FRDS1_repair oligo rv** | 17282 | TAGTAGCCTTTCGTCGTTCTTGCCAATGACTTGAGCCTTGA<br>CATTGATTCTAGCACCACCTTACAATTACGGATGAGGGGTT<br>GTTGAAGTGAAGATAAGAAAAAAAAATGGGGTGCTATG |
| **IDP2_repair oligo fw** | 17288 | GCTGCTCAGGCACGAGAATAGGAGGTAAGAAGGTAACGT<br>ACGTATATATATAAAATCGTAACTAAAGATTTGGCGCTCAT<br>TCTCGGTAAGTCTGAAAGATCCGCTTATGTTACTACCGAG |
| **IDP2_repair oligo rv** | 17289 | CTCGGTAGTAACATAAGCGGATCTTTCAGACTTACCGAGA<br>ATGAGCGCCAAATCTTTAGTTACGATTTTATATATATACGT<br>ACGTTACCTTCTTACCTCCTATTCTCGTGCCTGAGCAGC |
| **GPD1_repair oligo fw** | 17444 | GAAACAATTGTATATTGTACACCCCCCCCCTCCACAAACAC<br>AAATATTGATAATATAAAGATTTATTGGAGAAAGATAACA<br>TATCATACTTTCCCCCACTTTTTTCGAGGCTCTTCTATA |
| **GPD1_repair oligo rv** | 17445 | TATAGAAGAGCCTCGAAAAAAGTGGGGGAAAGTATGATA<br>TGTTATCTTTCTCCAATAAATCTTTATATTATCAATATTTGT<br>GTTTGTGGAGGGGGGGGGTGTACAATATACAATTGTTTC |
| **ALD3_repair oligo fw** | 17450 | TGATTTTATTTGTAAATAGTTATCAACGCCGGTGTCGCCTG<br>ATTCTCTACCAATACCACTTTTTCTTTTGGCTAATTTTCTAA<br>ATGTGTATAATCTATATCTCTATGATACAAGTCCAAG |

| | | |
|---|---|---|
| **ALD3_repair oligo rv** | 17451 | CTTGGACTTGTATCATAGAGATATAGATTATACACATTTAG AAAATTAGCCAAAAGAAAAAGTGGTATTGGTAGAGAATC AGGCGACACCGGCGTTGATAACTATTTACAAATAAAATCA |
| **GPP2_pURA3_fw** | 17752 | AACCAGCTAGTGTGTACCAGATCAGTGGAAAAACATAAAA CAATAAAAACAATATTCGGATGAGTATTTTCAATAAATTTG TAGAGGACT |
| **GPP2_tURA3_rv** | 17753 | CGAATATAGAATAGGACTGTATCTGAGAATTATTACTTAA ATATGTTCGATTTTAGAGGATTGCTTTTGTTCCACTACTTTT TG |
| **X2_pMPC3_fw** | 18025 | TCACAGAGGGATCCCGTTACCCATCTATGCTGAAGATTTAT CATACTATTCCTCCGCTCGTTATTTTGGTCGCCCCTTTAAC |
| **X2_tMPC3_rv** | 18026 | GTCATAACTCAATTTGCCTATTTCTTACGGCTTCTCATAAAA CGTCCCACACTATTCAGGATCCTGCTGCTAAAAAGGTC |

#### Table S8 List of primers used the verify removal or insertion of a gene.

| Primer name | Primer number | Sequence (5' to 3') |
|---|---|---|
| **AAC3 - CTRL FW** | 243 | GCTTCCAATGGCCTCCTCACCG |
| **AAC3 - CTRL RV** | 244 | GGGGGGGAATGCACTTAACAAAGAACC |
| **URA3_dg rev** | 1065 | CCCGGTAAAGCATTTCGAAG |
| **TKL2 dis500 fw** | 1360 | TCTTAATGGTGGCTCGCTGTC |
| **TKL2 dis500 rv** | 1361 | TCAATGCAGCCCATACACTC |
| **URA3_dg rev Inside** | 1741 | GAGCCCTTGCATGACAATTC |
| **GPD1_dg fw** | 2016 | CCCACCCACACCACCAATAC |
| **GPD1_dg rev** | 2017 | CGGACGCCAGATGCTAGAAG |
| **URA3_dg fw Inside** | 2891 | CATGGAGGGCACAGTTAAGC |
| **URA3_dg rev Insert** | 4905 | TCTTGGAAACGCTGCCCTAC |
| **OAC1_fw** | 6360 | AGTCAGGGTCTGCCGATATG |
| **OAC1_rv** | 6361 | CTTTCCGGTCCATGTTCGTC |
| **FW_gnd2KO_check** | 7258 | TCTGACAGGTGGCAGTTTCC |
| **RV_gnd2KO_check** | 7259 | ATCCGAAAGGCGGCAATAGG |
| **URA3_dg fw** | 7372 | GGGCAACGGTTCATCATCTCATGG |
| **FW_x-2_outside** | 7376 | GGTCTAGGCCTGCATAATCG |
| **RV_X-2_outside** | 7377 | TGCGGCATCATGTCTACTTG |
| **SHH3 dg fw** | 9450 | GTGACATGGCTCTTGCGTTC |
| **SHH3 dg rv** | 9451 | ACTTGCATCGCATCCAGTTC |
| **CTP1_dg fw** | 9493 | TGTCTGGGAACTCGGCTAAC |
| **CTP1_dg rv** | 9494 | TGGTTGGCTGTTTTTGTGGC |
| **GPP2_dg fw** | 9501 | AGTGTTCCGTCCGCGTCATC |
| **GPP2_dg rv** | 9502 | GGCGTTACACAGCCCAATCC |
| **SOL4_dg fw** | 9506 | GGGTGGACGTTTAAGCATAC |
| **SOL4_dg rv** | 9507 | GTATCACCGGGTGAGCTATG |
| **IDP1_dg_fwd** | 12305 | GAGAACCCGAGGACGATGTC |
| **IDP1_dg_rev** | 12306 | AACAGCCCCCTAAAACACGG |
| **IDP2_dg_fwd** | 12307 | GTGGGTTCTCACCGGGTTAG |
| **IDP2_dg_rev** | 12308 | ACTACCAGCTGGGATAGGGG |
| **PYC2_dg fw** | 12518 | CTGTGATTGGCAGAGAGGGG |
| **PYC2_dg rv** | 12519 | AATTCGTTTGCGCCAAGTCC |
| **SDH1b _dg fw** | 12525 | CTCCGGACTTATGTGGCTCC |
| **SDH1b _dg rv** | 12526 | ACTGAGGGAGAGGGTAAGG |
| **SHH4_dg fw** | 12533 | GCGCACTAGTTTGATGCCTG |
| **SHH4_dg rv** | 12534 | TGGCCTGCAAACTAAAACGC |
| **CIT3_dg fw** | 12540 | AGTGACACACCATGGTAGCG |
| **CIT3_dg rv** | 12541 | GACATCGGGGAAGTGGCTAC |
| **NQM1_dg fw** | 12572 | CCTTGATCTGGCTCTGGCTC |
| **NQM1_dg rv** | 12573 | CGCAAGGTAATTACGCCACG |
| **AAC1_dg fw** | 13823 | TTATCGGCGACTCAGCGTAC |
| **AAC1_dg rv** | 13824 | TTTGAGAGAGCACCGTCCAC |
| **SAL1_dg fw** | 13833 | TACCGGCCTTTCCAATACCC |
| **SAL1_dg rv** | 13834 | ATCTGGTTTGCGTTCGCATC |
| **ODC1_dg fw** | 13843 | AATGCGGCGACCAGAGATAG |
| **ODC1_dg rv** | 13844 | CCCCAGTCAAGTTGCATTGC |

| | | |
|---|---|---|
| **ODC2_dg fw** | 13849 | GTGCAGTCGCGATCACTTTC |
| **ODC2_dg rv** | 13850 | TCCCTGCGATCCGTTATGTG |
| **NDT2_dg fw** | 13856 | TTCTCGACATACTCGCGCTG |
| **NDT2_dg rv** | 13857 | CAGCTATTTCGGCAGTGCAC |
| **MPC3_dg fw** | 13862 | TGGTTTGGTCACGCAAAACC |
| **MPC3_dg rv** | 13863 | AAGTAGAAGAAGGCCGCGAC |
| **FRDS1_dg fw** | 17283 | TCGCAGGGTTTCTTCACAGG |
| **FRDS1_dg rv** | 17284 | ACAATCTTCAGGCCCGTCTG |
| **ALD3_dg fw** | 17448 | GCTTAGCTTACATGGCTGCG |
| **ALD3_dg rv** | 17449 | TTTAGGGAGGAGACAGGGAG |

#### Table S9 List of primers to construct and verify strain IMX1331

| Primer name | Primer number | Sequence (5' to 3') |
|---|---|---|
| ***Cas9* DNA fragment** | | |
| CAN1DelcassFW | 2873 | TCAGACTTCTTAACTCCTGTAAAAACAAAAAAAAAAAAAGGCATAG CAATAAGCTGGAGCTCATAGCTTC |
| A-CYC1t-rv | 4653 | GTGCCTATTGATGATCTGGCGGAATGTCTGCCGTGCCATAGCCATG CCTTCACATATAGTCCGCAAATTAAAGCCTTCGAG |
| **natNT2 DNA fragment** | | |
| tagA-pUG | 3093 | ACTATATGTGAAGGCATGGCTATGGCACGGCAGACATTCCGCCAGA TCATCAATAGGCACCTTCGTACGCTGCAGGTCGAC |
| CAN1 KO rv | 5542 | CTATGCTACAACATTCCAAAATTTGTCCCAAAAAGTCTTTGGTTCAT GATCTTCCCATACGCATAGGCCACTAGTGGATCTG |
| **Diagnostic primers to check integration** | | |
| **CAN1 cut rv** | 5829 | AGAAGAGTGGTTGCGAACAGAG |
| **m-PCR-HR4-RV** | 2673 | TGAAGTGGTACGGCGATGC |
| **m-PCR-HR2-FW** | 2668 | ACGCGTGTACGCATGTAAC |
| **Reverse primer pUGamds backbone** | 8442 | CGGGTGACCCGGCGGGGAC |
| **Nat Ctrl Fw** | 2620 | GCCGAGCAAATGCCTGCAAATC |
| **Can1RV** | 2615 | GAAATGGCGTGGGAATGTGA |

#### Table S10 Strain transformations

| Strain | Parental strain | Mutations | gRNA plasmid | Repair fragment |
|---|---|---|---|---|
| | | **Unreduced background** | | |
| **IMK588** | CEN.PK113-7D | *oac1Δ* | - | *KanMX* repair fragment |
| **IMK815** | IMX581 | *gnd2Δ tkl2Δ sol4Δ* | pUDR286 (*TKL2, SOL4*) + pUDR287 (*GND2*) | *SOL4*: 9504 + 9505<br>*TKL2*: 9509 + 9510<br>*GND2*: 7299 + 7300 |
| **IMX1592** | IMK815 | *nqm1Δ* | pUDR353 (*NQM1*) | *NQM1*: 12570 + 12571 |
| **IMX1694** | IMX581 | *pyc2Δ sdh1bΔ shh3Δ shh4Δ* | pUDR354 (*PYC2, SDH1b*) + pUDR355 (*SHH3, SHH4*) | *PYC2:* 12516 + 12517<br>*SDH1b:* 12523 + 12524<br>*SHH3:* 9448 + 9449<br>*SHH4*: 12531 + 12532 |
| **IMX1805** | IMX1694 | *cit3Δ* | pUDR351 | *CIT3:* 12538 + 12539 |
| **IMX2230** | IMX581 | *odc1Δ odc2Δ ndt2Δ ctp1Δ* | pUDR606 (*ODC1, ODC2*)<br>pUDR460 (*CTP1, NDT2*) | *ODC1*: 13841 + 13842<br>*ODC2*: 13847 + 13848<br>*NDT2*: 13854 + 13855<br>*CTP1*: 9491 + 9492 |
| **IMX2360** | IMX581 | *aac1Δ aac3Δ sal1Δ mpc3Δ* | pUDR458 (*AAC1, AAC3*)<br>pUDR462 (*SAL1, MPC3*) | *AAC1*: 13821 + 13822<br>*AAC3*: 13827 + 13828<br>*SAL1*: 13831 + 13832<br>*MPC3*: 13860 + 13861 |
| **IMX2391** | IMX581 | *odc2Δ* | pUDR688 (*ODC2*) | *ODC2*: 13847 + 13848 |
| **IMX2396** | IMX581 | *ctp1Δ odc1Δ odc2Δ* | pUDR686 (*ODC1, CTP1*)<br>pUDR688 (*ODC2*) | *ODC1*: 13841 + 13842<br>*ODC2*: 13847 + 13848<br>*CTP1*: 9491 + 9492 |
| **IMX2397** | IMX2360 | *odc2Δ* | pUDR688 (*ODC2*) | *ODC2*: 13847 + 13848 |
| **IMX2404** | IMX581 | *ctp1Δ odc1Δ ndt2Δ* | pUDR686 (*ODC1, CTP1*)<br>pUDR687 (*NDT2*) | *ODC1*: 13841 + 13842<br>*NDT2*: 13854 + 13855<br>*CTP1*: 9491 + 9492 |
| **IMX2408** | IMX2360 | *ctp1Δ odc1Δ ndt2Δ* | pUDR686 (*ODC1, CTP1*)<br>pUDR687 (*NDT2*) | *ODC1*: 13841 + 13842<br>*NDT2*: 13854 + 13855<br>*CTP1*: 9491 + 9492 |
| **IMX2416** | IMX2360 | *ctp1Δ odc1Δ odc2Δ* | pUDR686 (*ODC1, CTP1*)<br>pUDR688 (*ODC2*) | *ODC1*: 13841 + 13842<br>*ODC2*: 13847 + 13848<br>*CTP1*: 9491 + 9492 |
| **IMX2508** | IMX581 | *ctp1Δ* | pUDR738 (*CTP1*) | *CTP1*: 9491 + 9492 |
| **IMX2527** | IMX581 | *odc1Δ odc2Δ* | pUDR606 (*ODC1, ODC2*) | *ODC1*: 13841 + 13842<br>*ODC2*: 13847 + 13848 |
| **IMX2466** | IMX581 | *frds1Δ* | pUDR722 (*FRDS1*) | *FRDS1*: 17281 + 17282 |
| **IMX2467** | IMX581 | *idp1Δ* | pUDR723 (*IDP1*) | *IDP1*: 12295 + 12296 |
| **IMX2468** | IMX581 | *idp2Δ* | pUDR724 (*IDP2*) | *IDP2*: 17288 + 17289 |

| | | | | |
|---|---|---|---|---|
| **IMX2469** | IMX581 | *idp1Δ idp2Δ* | pUDR725 (*IDP1*, *IDP2*) | *IDP1*: 12295 + 12296<br>*IDP2*: 17288 + 17289 |
| **IMX2470** | IMX581 | *frds1Δ idp1Δ idp2Δ* | pUDR722 (*FRDS1*)<br>pUDR725 (*IDP1*, *IDP2*) | *FRDS1*: 17281 + 17282<br>*IDP1*: 12295 + 12296<br>*IDP2*: 17288 + 17289 |
| **IMX2509** | IMX581 | *ald3Δ gpd1Δ gpp2Δ* | pUDR739 (*ALD3*)<br>pUDR740 (*GPD1, GPP2*) | *ALD3*: 17450 + 17451<br>*GPD1*: 17444 + 17445<br>*GPP2*: 9499 + 9500 |
| **IMX2510** | IMX581 | *ald3Δ* | pUDR739 (*ALD3*) | *ALD3*: 17450 + 17451 |
| **IMX2512** | IMX581 | *ald3Δ gpd1Δ* | pUDR741 (*ALD3, GPD1*) | *ALD3*: 17450 + 17451<br>*GPD1*: 17444 + 17445 |
| **IMX2513** | IMX581 | *ald3Δ gpp2Δ* | pUDR742 (*ALD3, GPP2*) | *ALD3*: 17450 + 17451<br>*GPP2*: 9499 + 9500 |
| **IMX2612** | IMX581 | *gpd1Δ gpp2Δ* | pUDR740 (*GPD1, GPP2*) | *GPD1*: 17444 + 17445<br>*GPP2*: 9499 + 9500 |
| | | **Reduced background** | | |
| **IMK814** | IMX1331 | *gnd2Δ tkl2Δ sol4Δ* | pUDR286 (*TKL2, SOL4*) +<br>pUDR287 (*GND2*) | *SOL4*: 9504 + 9505<br>*TKL2*: 9509 + 9510<br>*GND2*: 7299 + 7300 |
| **IMX1591 (CCMin 1)** | IMK815 | *nqm1Δ* | pUDR353 (*NQM1*) | *NQM1*: 12570 + 12571 |
| **IMX1713** | IMX1591 | *pyc2Δ sdh1bΔ shh3Δ shh4Δ* | pUDR354 (*PYC2, SDH1b*) +<br>pUDR355 (*SHH3, SHH4*) | *PYC2:* 12516 + 12517<br>*SDH1b:* 12523 + 12524<br>*SHH3:* 9448 + 9449<br>*SHH4*: 12531 + 12532 |
| **IMX1806 (CCMin 2)** | IMX1713 | *cit3Δ* | pUDR351 | *CIT3:* 12538 + 12539 |
| **IMX1984** | IMX1806 | *aac1Δ aac3Δ sal1Δ mpc3Δ* | pUDR458 (*AAC1, AAC3*)<br>pUDR462 (*SAL1, MPC3*) | *AAC1*: 13821 + 13822<br>*AAC3*: 13827 + 13828<br>*SAL1*: 13831 + 13832<br>*MPC3*: 13860 + 13861 |
| **IMX2231** | IMX1984 | *odc1Δ odc2Δ ndt2Δ ctp1Δ* | pUDR606 (*ODC1, ODC2*)<br>pUDR460 (*CTP1, NDT2*) | *ODC1*: 13841 + 13842<br>*ODC2*: 13847 + 13848<br>*NDT2*: 13854 + 13855<br>*CTP1*: 9491 + 9492 |
| **IMX2394** | IMX1984 | *odc2Δ* | pUDR688 (*ODC2*) | *ODC2*: 13847 + 13848 |
| **IMX2405** | IMX1984 | *ctp1Δ odc1Δ* | pUDR686 (*ODC1, CTP1*) | *ODC1*: 13841 + 13842<br>*CTP1*: 9491 + 9492 |
| **IMX2406** | IMX1984 | *ctp1Δ odc1Δ odc2Δ* | pUDR686 (*ODC1, CTP1*)<br>pUDR688 (*ODC2*) | *ODC1*: 13841 + 13842<br>*ODC2*: 13847 + 13848<br>*CTP1*: 9491 + 9492 |
| **IMX2407 (CCMin 3)** | IMX1984 | *ctp1Δ odc1Δ ndt2Δ* | pUDR686 (*ODC1, CTP1*)<br>pUDR687 (*NDT2*) | *ODC1*: 13841 + 13842<br>*NDT2*: 13854 + 13855<br>*CTP1*: 9491 + 9492 |
| **IMX2471** | IMX2407 | *frds1Δ* | pUDR722 (*FRDS1*) | *FRDS1*: 17281 + 17282 |
| **IMX2472** | IMX2407 | *idp1Δ* | pUDR723 (*IDP1*) | *IDP1*: 12295 + 12296 |
| **IMX2473** | IMX2407 | *idp2Δ* | pUDR724 (*IDP2*) | *IDP2*: 17288 + 17289 |

| | | | | |
|---|---|---|---|---|
| **IMX2474** | IMX2407 | *idp1Δ idp2Δ* | pUDR725 (*IDP1*, *IDP2*) | *IDP1*: 12295 + 12296<br>*IDP2*: 17288 + 17289 |
| **IMX2475 (CCMin 4)** | IMX2407 | *frds1Δ idp1Δ idp2Δ* | pUDR722 (*FRDS1*)<br>pUDR725 (*IDP1*, *IDP2*) | *FRDS1*: 17281 + 17282<br>*IDP1*: 12295 + 12296<br>*IDP2*: 17288 + 17289 |
| **IMX2511** | IMX2475 | *ald3Δ* | pUDR739 (*ALD3*) | *ALD3*: 17450 + 17451 |
| **IMX2519 (CCMin 5)** | IMX2475 | *ald3Δ gpd1Δ gpp2Δ* | pUDR739 (*ALD3*)<br>pUDR740 (*GPD1, GPP2*) | *ALD3*: 17450 + 17451<br>*GPD1*: 17444 + 17445<br>*GPP2*: 9499 + 9500 |
| **IMX2520** | IMX2475 | *ald3Δ gpd1Δ* | pUDR741 (*ALD3, GPD1*) | *ALD3*: 17450 + 17451<br>*GPD1*: 17444 + 17445 |
| **IMX2521** | IMX2475 | *ald3Δ gpp2Δ* | pUDR742 (*ALD3, GPP2*) | *ALD3*: 17450 + 17451<br>*GPP2*: 9499 + 9500 |
| **IMX2538 (minimal CCM strain)** | IMX2520 | *gpp2Δ::URA3* | - | *URA3* repair fragment |
| **IMX2640** | IMX1984 | *X2::pMPC3-MPC3-tMPC3* | pUDR376 (*X2*) | *MPC3* repair fragment |
| **IMX2641** | IMX2519 | *X2::pMPC3-MPC3-tMPC3* | pUDR376 (*X2*) | *MPC3* repair fragment |
